# Probe-Based Identification of Metal-Binding Sites Using Deep Learning Representations

**DOI:** 10.1101/2025.10.04.680417

**Authors:** Shijie Xu, Akira Onoda

## Abstract

Metalloproteins play indispensable roles in a multitude of cellular processes. They incorporate metal ions as vital cofactors to catalyze biochemical reactions, stabilize protein structures, and mediate electron transfer. Given their prevalence and importance, identifying metal-binding sites within proteins remains a challenging task, due to the intricate complexity of the protein environments and the promiscuous binding behavior of metal ions. Although computational approaches for predicting metal-binding sites have been developed for decades, their performance often suffers from limited accuracy due to constrained methodologies and data scarcity. Here, we introduce PRIME, a hybrid deep learning framework that harnesses both evolutionary and structural signals to predict metal-binding sites with high accuracy and efficiency. PRIME employs protein language models (PLMs) and pre-trained structure models (PSMs), following the paradigm of deep representation learning, to extract essential information of protein sequences and structures. PRIME integrates a novel probe generation algorithm that bridges sequence- and structure-based predictions by efficiently scanning candidate binding sites. The resulting framework achieves superior accuracy across a wide range of metal ions, including both abundant ions such as Zn^**2+**^ and Ca^**2+**^ and less abundant ions such as K^**+**^ and Na^**+**^, surpassing the performance of existing methods. In addition, ablation analysis shows that PSMs significantly enhance the accuracy of metal-binding site prediction. Case studies on AlphaFold2 models, along with the high prediction speed of PRIME, further demonstrate its potential for high-throughput applications in metalloproteomics.

## 1 Main

Metalloproteins play essential roles in various cellular processes, including structural stabilization, catalysis, electron transfer, and signal transduction. They constitute nearly one-third of all deposited protein structures [1, 2] and exhibit an extraordinary diversity of functions. For example, a deficiency in metal ions can result in severe health issues and critical human diseases. Zinc-binding proteins involved in immune responses may lose functionality with age due to zinc deficiency [3] while iron-deficiency can lead to anemia, given iron’s vital role in hemoglobin synthesis [4]. Metalloenzymes also play important roles in industry, i.e., they utilize a protein scaffold to coordinate metal ions, thereby catalyzing reactions with desired stereo-selectivity and efficiency.

Importantly, metal binding is promiscuous, and certain ions may inhibit the binding of others. Cellular toxicity is the result of nonessential or toxic ions outcompeting the binding of essential ions. Calcium-binding proteins function as sensitive detectors of environmental calcium concentrations [5, 6] and regulators of downstream proteins [7]. The selective binding of K^+^ ions to proteins enables the formation of ion channels through protein tetramerization [8], which is crucial for ion transport. Hg^2+^ ions exhibit pronounced toxicity, and mercury-resistant proteins such as MerR [9] have evolved to bind mercury with exceptional sensitivity, while displaying comparatively lower affinities for Cd^2+^ and Zn^2+^ [10]. Therefore, predicting the specificity of metal binding remains a significant challenge.

Identifying binding residues out of whole protein residues is also challenging. Although some ions, such as zinc, have strong preferences for specific chemical groups (e.g., His or Cys), many binding sites also involve less specific interactions with atoms such as backbone carbonyl oxygen atoms. For instance, alkaline earth metal ions such as Ca^2+^ preferentially bind regions enriched in negatively-charged residues including Asp and Glu. Furthermore, the dynamic nature of proteins complicates the reliability of deposited structures, as interactions between metal ions and protein residues can induce conformational changes [7, 9, 11]. Conventional computational approaches, including molecular dynamics simulations, encounter difficulties in accurately modeling transition metal interactions [12], as most force fields are optimized for organic molecules or biomacromolecules [13]. Hybrid quantum mechanics/molecular mechanics (QM/MM) methods provide improved accuracy. However, these are computationally demanding and constrained in both timescale and system size [14].

While recent advances in structure prediction programs such as the AlphaFold series [15] and RosettaFold [16] have enabled the generation of high-quality protein structures, these models often omit metal-binding sites. On the other hand, AlphaFold3 can predict ligand-binding sites if ions are explicitly included as input. However, this step is often omitted in practice. In addition to computational predictions, experimental techniques such as cryo-EM provide structural information, but achieving resolution sufficient to resolve metal-binding sites remains challenging [17]. As a result, computational approaches for predicting metal-binding sites are essential, particularly for less abundant metal ions that are underrepresented in experimental datasets.

Numerous computational strategies have been devised to predict metal-binding proteins. Template-based approaches such as MIB [18] and MIB2 [19] depend on 2 know structural motifs and are constrained by template availability. BioMetAll [20] constructs probe grids around proteins and identify potential binding sites through geometric screening. Metal3D [12] is a deep learning framework that predicts metal-binding sites from protein structures and shows superior performance compared to BioMetAll and MIB/MIB2. On the other hand, sequence-based methods such as mebipred [21] leverage machine learning to identify proteins as metalloproteins at the protein level. The advent of protein language models (PLMs) has led to PLM-based methods such as LMetalSite [22] and M-Ionic [23], which are pre-trained on massive sequence databases (e.g., UniProt [24]) and extract residue-level features for metal site prediction. While PLM-based methods offer high speed and accuracy, they are limited by their reliance on sequence alone and lack the ability to model spatial geometry. ESMBind [25] attempts to bridge this gap by integrating PLM-derived features with structural energy minimization, but this system operates under the assumption that each protein contains only one binding site. SuperMetal [26], another recently developed deep learning model, predicts zinc-binding sites by generating candidate regions via diffusion processes and refining predictions with a second-stage predictor. Its use of graph neural networks (GNNs) [27] to interpret PLM features may obscure sequential patterns. Geometric information is not sufficient to predict when oligomeric proteins are absent in the structures (e.g., the AlphaFold2 models), because binding sites involving multiple binding partners may not be captured by methods which only analyze structures, due to the missing symmetrical mates. Such oligomeric proteins might have sufficient conservation to be detectable at the sequence level [28]. Other methods such as AlphaFill [29], AlphaFold3 [30], and Metal-installer [31] can predict with a prior knowledge of the existence of metal-binding sites, which is not in line with to the objective of the present study. Recent efforts [12, 32] have employed deep learning, particularly convolutional neural networks (CNNs) [33], to predict metal-binding sites from protein structures, yet their performance remains limited, especially for less abundant metal ions.

To address the limitations of deep learning methods that rely solely on either sequence or structural data, we propose PRIME (<u>PR</u>obe-based <u>I</u>dentification of <u>ME</u>talbinding sites using deep learning representations), a novel approach designed to enhance the accuracy of metal-binding sites prediction of proteins. PRIME introduces several key innovations by 1) jointly utilizing both sequence and structural information to improve both accuracy and efficiency of metal-binding prediction; 2) incorporating pre-trained deep representations of protein structures, allowing it to better capture structural patterns; and 3) including a novel probe generation algorithm to efficiently and accurately generate candidate binding sites. These innovations enable PRIME to achieve superior accuracy and computational efficiency across a wide range of metal ions, outperforming existing methods.

## 2 Results and discussion

### 2.1 PRIME

PRIME takes protein sequences and structures as inputs and predicts the metal-binding sites in proteins (Figure 1). A sequence-based predictor, (PRIME-seq), ranks the propensity of residue-level binding by using learning motifs from sequence data, aided by high-dimensional representations extracted through protein language models (PLMs). Residues with scores above a certain threshold are selected, and structural context around them is used to generate candidate binding sites, which are referred to as “probes”. This terminology, borrowed from BioMetAll [20], denotes physically plausible regions around the selected residues. The purpose of probe generation is to ensure comprehensive coverage of true binding sites while excluding regions less likely to contain binding sites. These probes are then refined through a structure-based predictor (PRIME-probe), followed by additional post-processing steps to yield the final predictions.

**Fig. 1.**
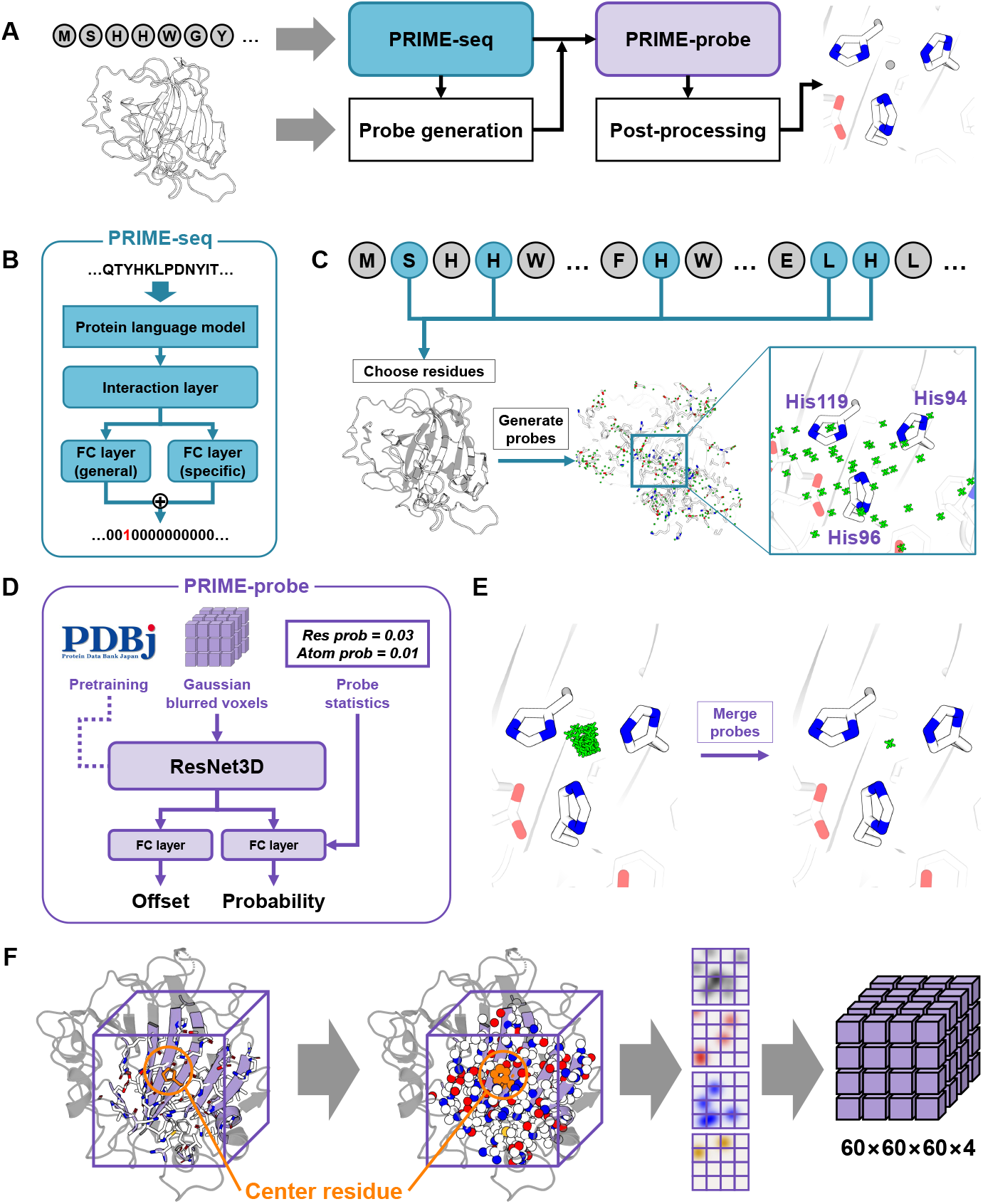
(A) The overall architecture of PRIME consists of two deep learning modules, PRIME-seq and PRIME-probe, alongside a probe generation algorithm and a series of post-processing procedures. PRIME accepts protein sequences and structures as input, predicting metal-binding sites of proteins. (B) The architecture of PRIME-seq. A protein language model (PLM) encodes the protein sequence into high-dimensional representations, which are processed by an interaction layer, such as a recurrent neural network (RNN) or transformer encoder, to capture residue interactions. Two fully connected (FC) layers follow: one is trained on all metal ions to model general binding propensities, and the other is trained for individual metal ions to capture specific binding patterns. The final predicted propensity is the sum of these outputs. (C) A probe generation algorithm utilizes the predictions from PRIME-seq to generate candidate binding sites (probes), taking into account both the geometrical context and predicted binding propensity of the residues. (D) The architecture of PRIME-probe. The local environments surrounding the probes are extracted from the protein structures and blurred using a Gaussian kernel to construct a voxel grid. This grid is then fed into a three-dimensional residual network (ResNet3D), which includes two FC layers, to predict both the offset vector and the binding probability of each probe. The offset vector is used to refine the probe position. Statistical features of the probes are also adopted as an input of PRIME-probe. To improve the accuracy of prediction, the ResNet3D is pretrained via a masked language modeling task, offering a better initialization for fine-tuning on metal-binding site prediction. (E) The refined probes are subsequently filtered based on their predicted probabilities and clustered to to eliminate spatial redundancy among probes. (F) Workflow of voxelization and Gaussian blurring. A residue is selected, and neighboring atoms within a cubic region of 10 Å centered at its alpha carbon are extracted from the protein structure. Only carbon, nitrogen, oxygen, and sulfur atoms are considered and blurred using a Gaussian kernel to produce a voxel grid.

PRIME attempts to leverage both sequence and structural information of proteins. This approach stems from the observation that metal-binding regions, particularly those coordinating transition metals, often comprise distinctive amino acid motifs discernible from sequence data. Moreover, when structural models fail to capture interactions between protein chains, these conserved sequence motifs can compensate for the missing structural information.

Recent advances in protein language models have revealed that deep learning representations of protein sequences are sufficiently powerful to capture sequence motifs [34–39], yet they do not process structural information. Structure-based approaches, such as BioMetAll and the MIB series, are often hindered by a high number of false positive predictions. While recent methods such as Metal3D have demonstrated notable progress in deep-learning-based metal-binding prediction, precision remains constrained, and analyses which include less abundant metal ions are less reliable [12, 40] because deep learning methods frequently suffer from limited training sets. An additional and pressing shortcoming of current structure-based methods lies in their inefficiency (e.g., BioMetAll), particularly with larger proteins, which hinders their application in high-throughput screening of metal-binding sites.

PRIME utilizes an efficient probe generation algorithm to integrate sequence and structural information, accurately identifying candidate binding sites within protein structures. Transition metal ions exhibit pronounced affinities for specific amino acids (Supplementary Fig. S1, S2, S3), a feature leveraged by PRIME to achieve high precision in binding site prediction. PRIME achieves over 99% coverage of binding sites for transition metal ions in the validation sets. Across the entire datasets of transition metals, PRIME missed only 8 out of 4193 binding sites for Zn^2+^, 2 out of 961 binding sites for Mn^2+^, 1 out of 325 binding sites for Cu^2+^, and 1 out of 346 binding sites for Co^2+^, with no missing sites for others. These analyses used less than 300 probes for each protein. For non-transition metals, PRIME missed only 6 out of 3359 binding sites for Ca^2+^ and 1 out of 2439 binding sites for Mg^2+^ and no missing sites for K^+^ and Na^+^, with approximately 1000 probes per protein (Supplementary Fig. S4). By focusing on the most probable binding regions, this strategy enhances the accuracy and efficiency of the structure-based predictor. Additionally, PRIME is trained and applied individually for each metal ion, rather than jointly across all ions [32, 40], enabling the model to capture the distinct binding patterns of each metal ion, which is essential given the frequent competition among metal ions for binding sites.

Self-supervised pretraining on PDB structures was found to significantly improve prediction accuracy and stability, particularly for larger models (Supplementary Fig. S5). The structure-based predictor (PRIME-probe) was pretrained using a masked language modeling task on 16,839 non-redundant PDB structures. A cubic region of 10 Å centered on the alpha carbon atom of a randomly selected residue was extracted, with the atoms of the selected residue masked. A three-dimensional residual network (ResNet3D), the 3D counterpart of the residual network [41], was employed to predict the masked residue types. Pretraining revealed a scaling law with respect to model size, demonstrating that larger models achieve superior performance on the masked language modeling task (Supplementary Fig. S6). The pretrained models were subsequently fine-tuned on the metal-binding prediction task, and the best model was selected based on validation performance.

#### 2.1.1 Metal-binding residue prediction

We compare several state-of-the-art predictors for metal-binding residues, including PRIME-seq, LMetalSite, M-Ionic, and MetalNet. The performance of these predictors for four abundant metal ions, Zn^2+^, Ca^2+^, Mg^2+^, and Mn^2+^, are illustrated in Figure 2 and Supplementary Fig. S7, through both precision-recall (PR) and receiver operating characteristic (ROC) curves. The first three methods, PRIME-seq, LMetal-Site, and M-Ionic, are all based on protein language models (PLMs), employing ESM [36] (PRIME-seq), ProtTrans [34] (LMetalSite), or a combination of both (M-Ionic). Their PR and ROC curves are similar, indicating that further improvement of PLM-based methods will be challenging under the current data size and comparable model architectures. In contrast, MetalNet employs a different approach, utilizing multiple sequence alignments (MSAs) to extract co-evolutionary signals and predict residue pairs involved in the same binding sites, indicating that pair-based prediction is less effective than residue-level prediction for metal-binding sites.

**Fig. 2.**
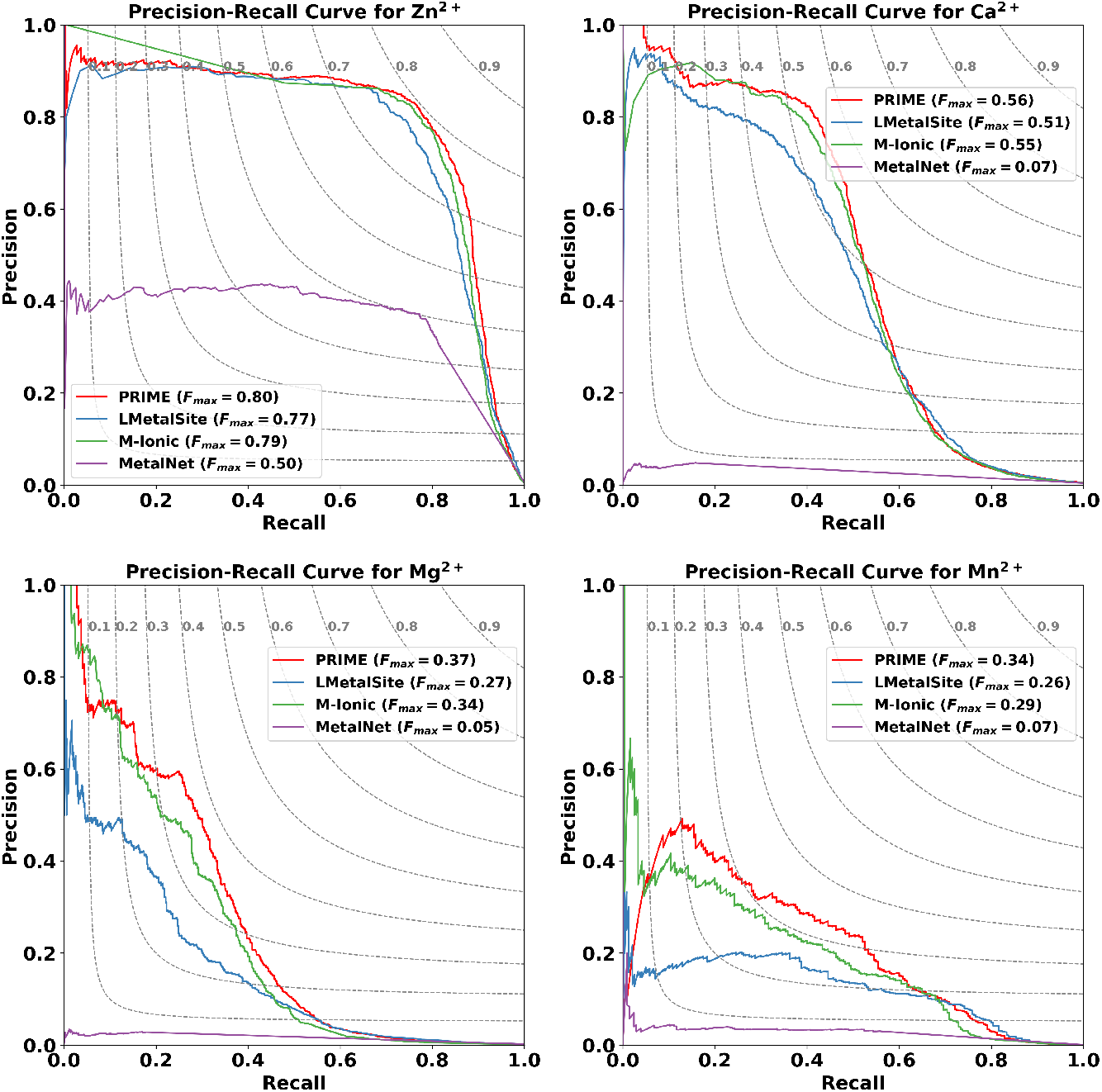
Precision-recall curves for the prediction of metal-binding residues by PRIME and other sequence-based methods for Zn^2+^, Ca^2+^, Mg^2+^, and Mn^2+^. The legends indicate the maximum F1 scores across all thresholds.

The prediction performance varies considerably among different metal ions, reflecting a positive correlation with the size of the training data set. As the most represented ion in the dataset, Zn^2+^ yields the highest accuracy. Ca^2+^ and Mg^2+^, non-transition metals whose binding is largely dominated by electrostatic forces, pose greater difficulty than Zn^2+^. Despite Mn^2+^ being a transition metal with clear residue preferences (Supplementary Fig. S3), its prediction remains challenging without structural information, indicating that sequence-based approaches are severely limited by data scarcity.

The limitations imposed by data scarcity become even more apparent when predicting a broader range of metal ions. For less abundant ions such as Fe^2+^, Co^2+^, Cu^2+^, K^+^, Ni^2+^, Cd^2+^, Mn^3+^, and Hg^2+^, PRIME-seq often yields an F1 score of zero, indicating a complete lack of non-zero precision or recall (Supplementary Table S1, Fig. S8). Given the limited availability of training data for these ions and the challenges in expanding these datasets in the near future, sequence-based methods alone appear insufficient. Nevertheless, these metals frequently exhibit high test AUC scores, suggesting that PRIME-seq still serves effectively in ranking candidate residues [42].

#### 2.1.2 Reduced search space

A key innovation of PRIME lies in its probe generation algorithm, which substantially reduces the search space within the atomic environment of a protein. To quantitatively assess the algorithm’s effectiveness, the ratio of the volume occupied by the generated probes to the total volume of the search space of heavy atoms (C/N/O/S) in the protein structure is computed. Monte Carlo integration is employed to estimate the volume, defined as the union of the van der Waals spheres of probes or the heavy atoms in the protein structure. The results reveal that most ions exhibit a markedly reduced search space, typically below 10% (Figure 3), whereas K^+^ ions display a relatively higher ratio, 46.26%, followed by Na^+^ at 12.47% and Ca^2+^ at 11.53%, due to their less binding specificities to residues. Overall, the probe generation algorithm of PRIME effectively minimizes the search space for metal-binding sites while maintaining high coverage of true binding sites (Supplementary Table S2 and Table S3).

**Fig. 3.**
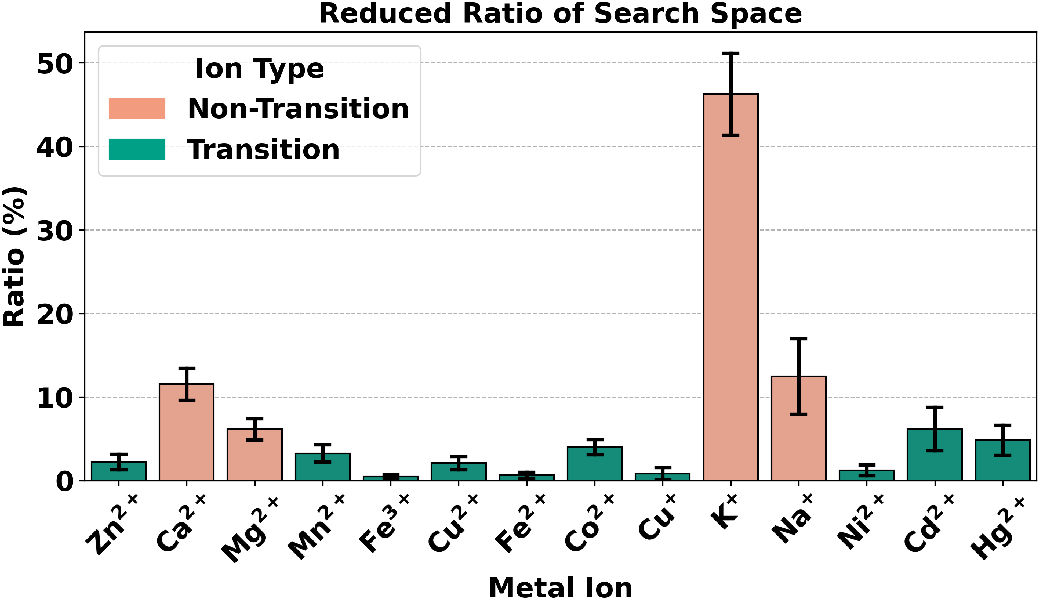
The reduced search space ratio of the probe generation algorithm. Certain alkali and alkaline earth metals, such as K^+^, Na^+^, and Ca^2+^, exhibit significantly higher ratios compared to transition metals, potentially due to their less specific binding residues.

#### 2.1.3 Metal-binding site prediction

A comparative analysis of structure-based metal-binding site prediction methods, including PRIME, BioMetAll, Metal3D, and AllMetal3D, was conducted on the zinc-binding test set (Figure 4A). PRIME attains an F1 score of 0.8920, surpassing all other methods, with AllMetal3D and Metal3D achieving 0.8072 and 0.7708, respectively, and BioMetAll yielding 0.6297. PRIME also achieves the second highest recall (0.9129) and precision (0.8920) among the methods. BioMetAll attains the highest recall (0.9828) but only 0.4633 precision, indicating that it identifies nearly all binding sites but with a substantial number of false positives. In contrast, PRIME achieves a balanced trade-off between recall and precision. AllMetal3D and Metal3D employ similar model architectures yet display different performance, suggesting that training strategies, such as joint training across metal ions in AllMetal3D, may influence outcomes. Additionally, PRIME achieves a comparable RMSE of 0.8900 Å, which is slightly higher than AllMetal3D (0.6703 Å) and Metal3D (0.5484 Å), but markedly lower than BioMetAll (1.1500 Å). This result is reasonable, as PRIME is optimized primarily for classification accuracy, with the distance loss down weighted by a factor of 0.1 (section 4.3.5). Consequently, PRIME demonstrates superior classification accuracy relative to positional accuracy.

**Fig. 4.**
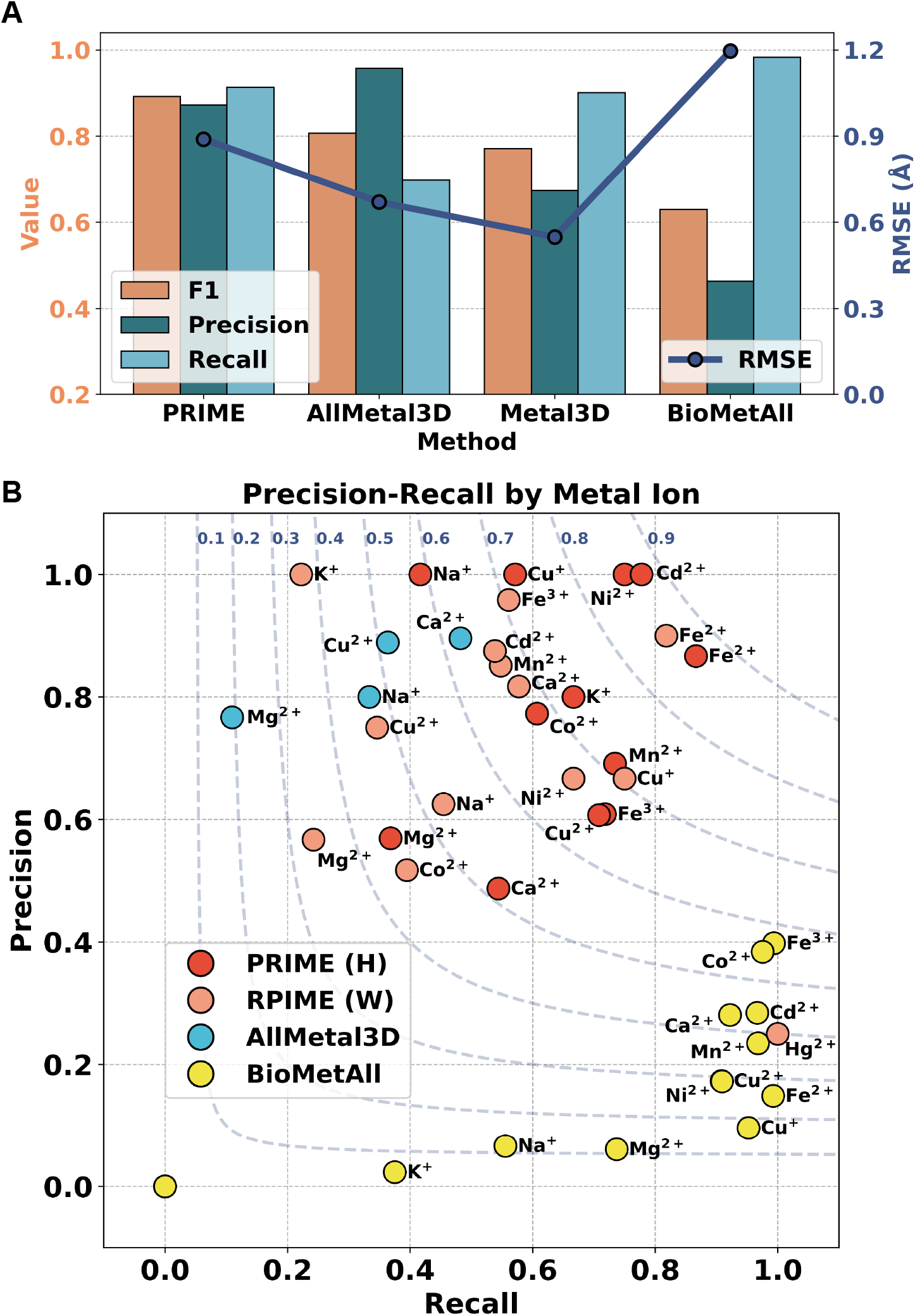
(A) Comparison of precision, recall, and F1 scores (left axis) and RMSE (right axis) for zinc-binding site prediction by PRIME and other structure-based methods. (B) Scatter plot of recall and precision for different methods on additional _11_metal ions, including Ca^2+^, Mg^2+^, Mn^2+^, Fe^3+^, Cu^2+^, Fe^2+^, Co^2+^, Cu^+^, K^+^, Na^+^, Ni^2+^, Cd^2+^, and Hg^2+^. PRIME (H) denotes PRIME trained with the hard mining strategy, and PRIME (W) denotes PRIME trained with the weighting strategy. Methods are distinguished by color. Isometric F1 score lines are indicated by blue dashed curves.

The performance of PRIME was further evaluated on additional metal ions, including Ca^2+^, Mg^2+^, K^+^, Na^+^, Fe^2+^, Ni^2+^, Cu^2+^, Co^2+^, Cd^2+^, and Hg^2+^, demonstrating consistently superior results across all ions (Figure 4B). Two PRIME variants, Hard Mining (H) and Weighting (W), display distinct performance profiles. The hard mining strategy emphasizes challenging negative samples, omitting easier cases, which is particularly effective for zinc-binding sites with high residue specificity. The weighting strategy incorporates all negative samples, promoting broader recognition of non-binding residues, and may be more suitable for metal ions with lower residue specificity, such as Ca^2+^. Among all metal ions, PRIME attains the highest F1 scores compared to other methods, followed by AllMetal3D and BioMetAll. AllMetal3D achieves higher precision but lower recall, while BioMetAll achieves higher recall but very low precision. Furthermore, AllMetal3D yields zero recall and precision for Mn^2+^ and Fe^3+^ ions, despite sufficient test data (Mn^2+^: 78, Fe^3+^: 45). This limitation arises from the joint training strategy employed by AllMetal3D, which is primarily trained on zinc, calcium, and magnesium sites, resulting in a preference for common metal ions and greatly limiting its performance on others.

### 2.2 Prediction on Cryo-EM and AlphaFold2 structures

Recent advances in experimental and computational methods, including cryogenic electron microscopy (cryo-EM) [43] and AlphaFold2 [15], have greatly expanded the repertoire of available protein structures. However, the limited resolution of cryo-EM structures often precludes precise identification of metal-binding sites [44]. Inaccuracies in AlphaFold2 models may compromise the reliability of computational predictions, including molecular docking [45]. Assessing the performance of PRIME on proteins lacking high-resolution crystallographic data is therefore crucial.

PRIME was evaluated on the human zinc-activated channel protein (hZAC, PDB ID: 8YX6), recently resolved by cryo-EM, which contains two zinc-binding sites per monomer [46]. PRIME accurately predicts 9 out of 10 experimentally determined Zn^2+^ binding sites, with site 1 and site 3 (Figure 5A, 5B) assigned high probabilities of 0.84 and 0.59, respectively. Additionally, PRIME identifies a site 2 with a probability of 0.63, which has not been reported or discussed in the literature and warrants further experimental validation. AllMetal3D was also applied to the same structure. Despite being trained predominantly on zinc-binding datasets, AllMetal3D predicts only two zinc binding sites, site 1 in chain A (0.54) and site 3 in chain B (0.61).

**Fig. 5.**
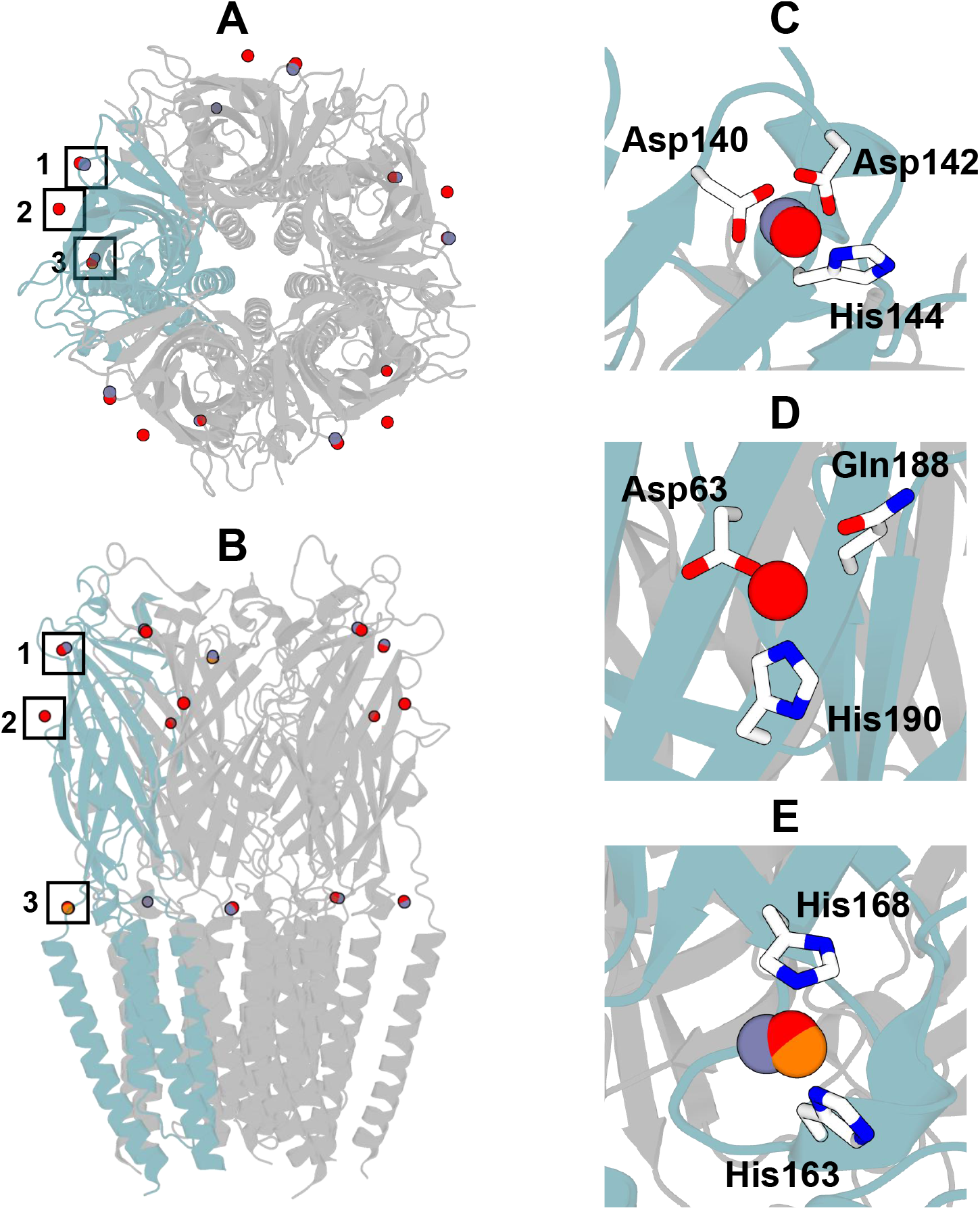
Prediction of zinc-binding sites in the human zinc-activated channel protein (hZAC). (A) Top view of the hZAC structure and predicted sites. (B) Front view of the hZAC structure and predicted sites. (C) Predicted binding site 1 on chain B. (D) Predicted binding site 2 on chain B. (E) Predicted binding site 3 on chain B. Chain A is shown in light blue, actual binding ions are shown in blue, predicted ions from PRIME are shown in red, and predicted ions from AllMetal3D are shown in orange.

Metallothioneins from *Arabidopsis thaliana* (UniProt ID: P43392) are small, highly conserved, cysteine-rich proteins that bind a variety of metal ions, functioning as chelators, mitigating Cu and Cd toxicity, and contributing to metal homeostasis [47]. As no crystallographic structures of metallothioneins are available, the AlphaFold2 model was employed for prediction. PRIME identifies plausible binding sites for Zn, Cu, and Cd based on the AlphaFold2 structure (Figure 6). Although experimental evidence for Hg and Ni binding is absent, homologous proteins from mouse (gi: BC036990) have been reported to bind Hg^2+^ and Ni^2+^ with high affinity [48, 49]. While AllMetal3D supports predictions for Cu^+^ and Cu^2+^, it does not identify any binding sites in this protein.

**Fig. 6.**
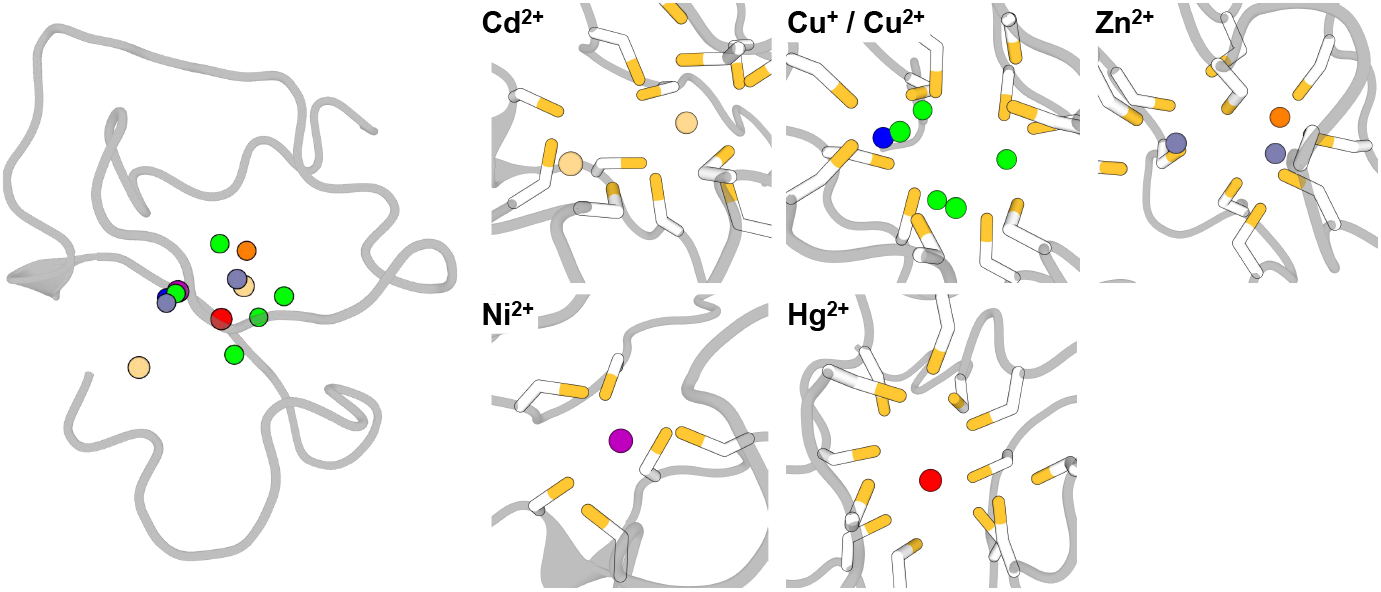
Predicted binding sites of metallothioneins for various metal ions and detailed views of each binding site, including Cd^2+^, Cu^+^/Cu^2+^, Zn^2+^, Ni^2+^, and Hg^2+^. Cu^+^ and Cu^2+^ are shown in green and blue, respectively. Zn^2+^ predictions from PRIME are colored orange, and those from AllMetal3D are blue. Coordinating cysteines are depicted as sticks.

### 2.3 Prediction of ion channels

Potassium channel KcsA-Fab from *Streptomyces lividans* (PDB ID: 1K4C) is a well-characterized tetrameric ion channel exhibiting selective affinity for K^+^ ions [8]. Its architecture is highly symmetrical, with four identical subunits forming a central channel. Each subunit contains a selectivity filter that facilitates the passage of K^+^ ions while excluding others. The K^+^ binding sites are located at the core of the filter (Figure 7A, 7B, 7C), where the ions are coordinated by oxygen atoms from the protein backbone. The geometry of the ion channel and the predicted metal-binding sites are shown in Figure 7.

**Fig. 7.**
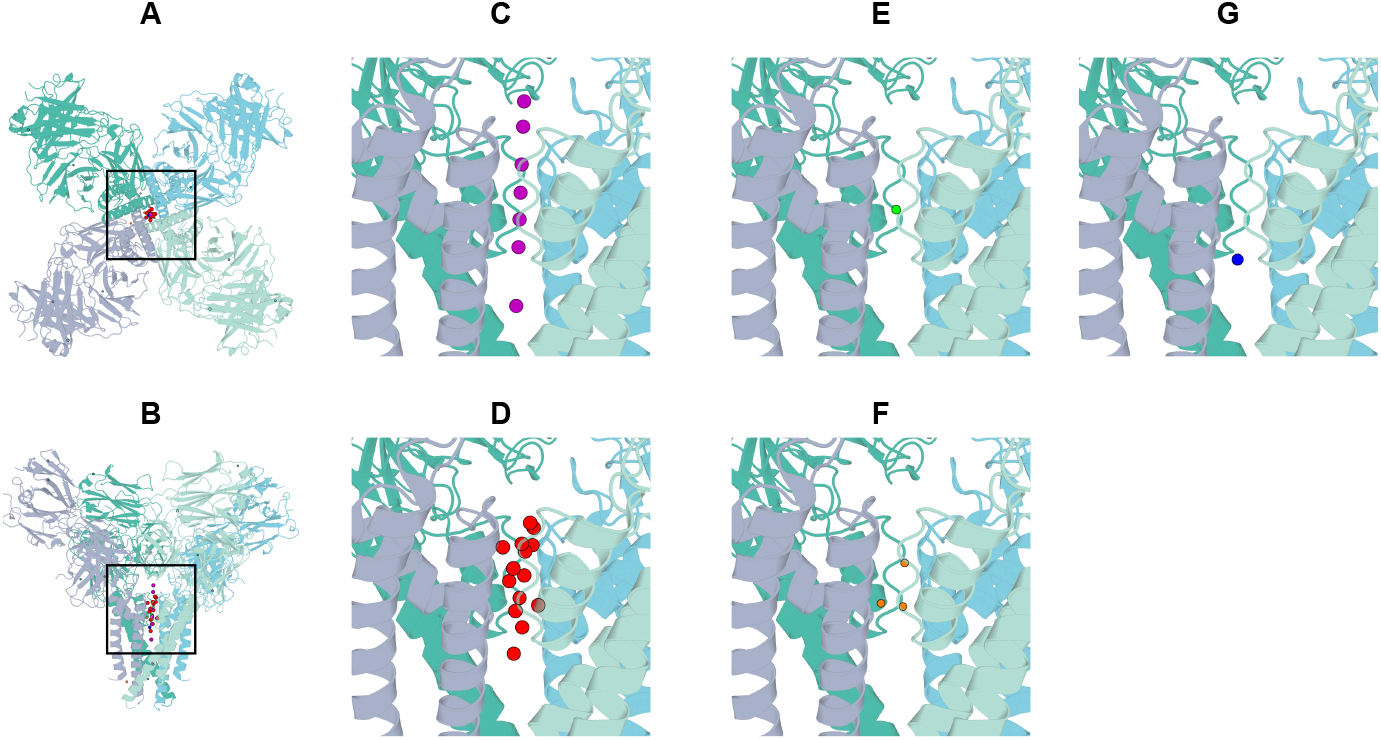
Tetrameric structure of the K^+^ ion channel in protein 1K4C. Each subunit is rendered in a distinct color. Symmetrical mates were generated using PyMOL, and the clustering distance in PRIME was set to 1.0 Å (default: 3.0 Å). Purple spheres indicate the experimentally determined K^+^ ions in the PDB structure, and red spheres denote the predictions by PRIME. (A) Top view of the tetrameric K^+^ channel. (B) Front view of the tetrameric K^+^ channel, with the ion pathway centered. (C) Enlarged view of the actual K^+^ ion in the structure. (D) Prediction of K^+^ ions, (E) Ca^2+^ ion, (F) Cd^2+^ ions, and (G) Na^+^ ion.

PRIME accurately predicts the K^+^ binding sites in the tetrameric structure (Figure 7D), revealing a series of binding sites extending along the channel. Additionally, predicted binding sites for Ca^2+^ and Na^+^ are identified (Figure 7E, 7G), with probabilities of 0.98 and 0.55, respectively. In contrast to the extensive distribution of K^+^ binding sites within the channel, Ca^2+^ and Na^+^ binding sites are limited to a single location, suggesting weaker binding affinities. Notably, PRIME does not predict any Mg^2+^ binding sites but identifies four Cd^2+^ binding sites with probabilities of 0.94, 0.91, 0.79, respectively, as shown in Figure 7F.

### 2.4 Prediction of apo and holo forms

Calmodulin predictions were conducted for both the apo form (PDB ID: 1CFD) and holo form (PDB ID: 1CLL), as illustrated in Figure 8. The ChimeraX [50] `morph` command generated ten interpolated transition states between the apo and holo forms, and calcium-binding sites were predicted for each state. In the early transition states (State 0 to State 5), no calcium ions were identified, while the EF-hand motifs progressively revealed binding sites in States 7 to 10. The predictions from PRIME correspond to the conformational changes and demonstrate robust performance. In the final state (1CLL), three binding sites were predicted rather than four, as observed in the crystal structure. This is apparently attributable to the pronounced flexibility of the helix-loop-helix regions.

**Fig. 8.**
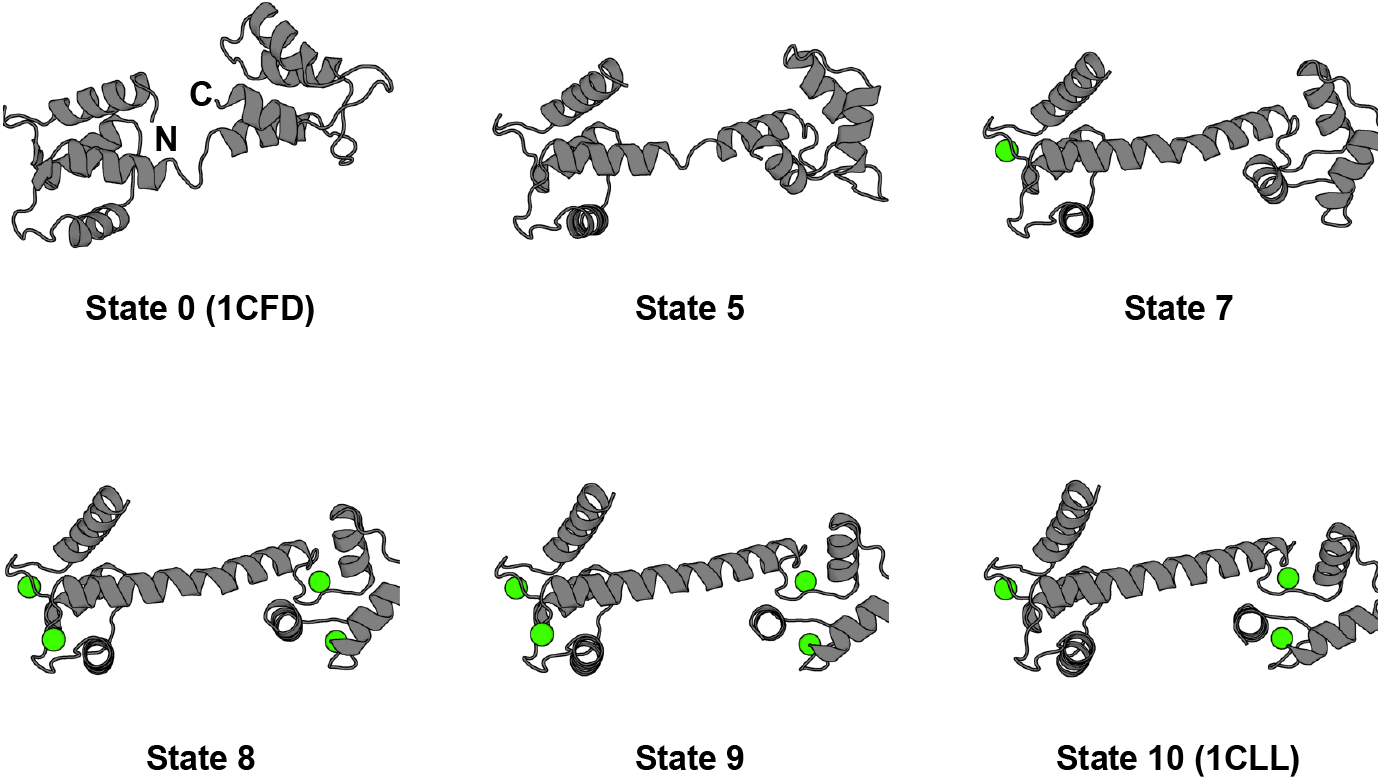
Prediction of calcium-binding sites in calmodulin by PRIME. Ten transition states were uniformly interpolated, with selected states shown as States 0, 5, 7, 8, and 10. State 0 corresponds to the apo form (PDB ID: 1CFD), and State 10 corresponds to the holo form (PDB ID: 1CLL). Predicted calcium-binding sites are depicted in green.

### 2.5 Prediction of less abundant metal ions

Benchmarking reveals that several less abundant metal ions, including Ni^2+^ and Cd^2+^, achieve notably high F1 scores despite limited training data (Figure 4C). To validate these results, three representative examples are presented. The SBP FpvC protein from *P. aeruginosa* (PDB ID: 6R3Z) is a nickel-binding protein featuring a coordination motif containing six His residues, which is rare among metalloproteins. PRIME accurately identifies this motif, with only minor spatial deviation from the annotated site (Figure 9A). The prediction for *Helicobacter pylori* UreE protein (PDB ID: 3TJ8), which mediates nickel insertion essential for urease activity, is also highlighted. The binding site is located at the interface of two subunits. PRIME yields two predictions, one for the monomer and one including the symmetrical mate (Figure 9B). The experimentally determined nickel ion (green) is coordinated by three histidine residues from one subunit and an aspartate from the other. PRIME predicts a single binding site with slight deviation in the monomeric model, and two closely positioned sites when the symmetrical mate is included, potentially reflecting the dynamic nature of nickel insertion. *Candidatus liberibacter asiaticus* ZnuA2 coordinates Cd^2+^ at a site of high specificity, functioning as a heavy metal exporter. PRIME accurately predicts the binding position, with the ion coordinated in a canonical octahedral geometry (Figure 9C).

**Fig. 9.**
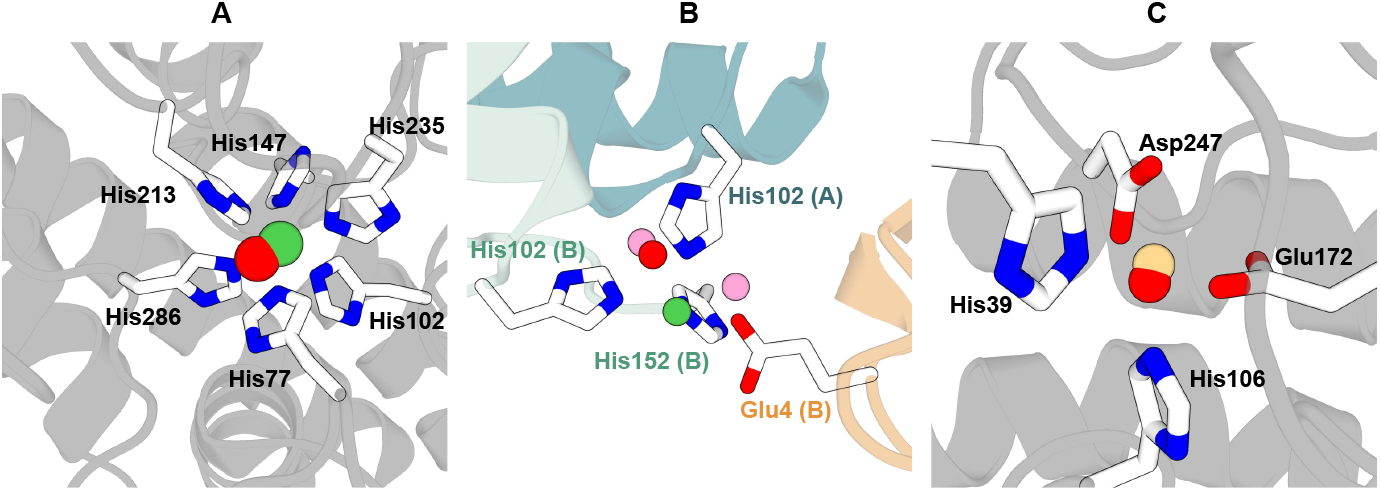
Prediction of Ni^2+^ and Cd^2+^ binding sites by PRIME. (A) Ni^2+^ binding site in SBP FpvC protein (PDB ID: 6R3Z). The green sphere indicates the location of the experimentally determined Ni^2+^ ion, and the red sphere indicates the PRIME prediction. (B) Ni^2+^ binding site in *Helicobacter pylori* UreE protein (PDB ID: 3TJ8). The green sphere indicates the location of the experimentally determined Ni^2+^ ion, the red sphere indicates the PRIME prediction for the monomer, and the pink sphere represents the prediction for the dimer. (C) Cd^2+^ binding site prediction in *Candidatus liberibacter asiaticus* ZnuA2 (PDB ID: 6IXI). The green sphere indicates the location of the experimentally determined Cd^2+^ ion, and the red spheres indicate the PRIME predictions.

### 2.6 Metal selectivity

The mercury-binding protein MerR acts as a transcriptional repressor-activator, exhibiting remarkable sensitivity and selectivity for Hg^2+^ ions. MerR has an exceptionally high association constant for Hg^2+^ at approximately 10^47^ M^−1^ [9]. It also binds Cd^2+^ and Zn^2+^ with association constants which are two to three orders of magnitude lower than that for Hg^2+^ [10]. PRIME predictions for the binding sites of MerR (PDB ID: 5CRL) are shown in Figure 10A, where all predicted sites coincide with the experimentally determined binding site, except for a Cd^2+^ site which has a probability of 0.79 of being located externally. In the left binding site, which contains more predicted metal ions, PRIME identifies binding sites with descending probabilities: Hg^2+^ (0.98), Zn^2+^ (0.95), and Cd^2+^ (0.79) (Figure 10B). Hg^2+^ is predicted with the highest probability, consistent with MerR’s selectivity for Hg^2+^ ions. The high probabilities for Zn^2+^ and Cd^2+^ correspond to their experimentally reported affinities, which are slightly lower than that of Hg^2+^. The reduced probabilities for Cu^2+^ and other ions may reflect structural or geometric constraints that limit MerR’s compatibility with these ions.

**Fig. 10.**
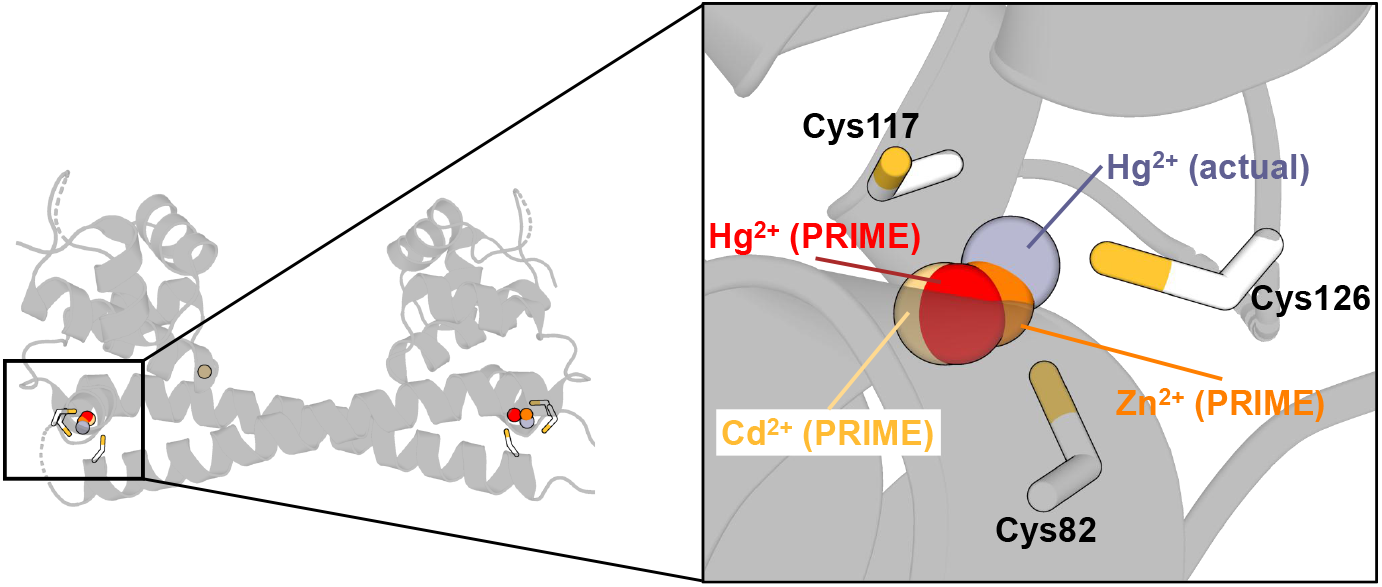
(A) Predicted metal-binding sites of MerR (PDB ID: 5CRL) by PRIME. Binding residues in two sites are shown as stick representations. (B) A zoomed view of the left binding site, displaying predicted metal ions: Hg^2+^ (red), Zn^2+^ (orange), Cd^2+^ (yellow), and the actual Hg^2+^ (gray) from the PDB structure.

### 2.7 Specificity of different metal ions

Proteins exhibit diverse binding preferences for metals, and a binding site can often accommodate diverse metal ions. For instance, Zn^2+^ and Co^2+^ share a similar binding residue distribution and are sometimes exchangeable in certain binding sites, while Zn^2+^ and Ca^2+^ are quite different. To investigate the specificity of PRIME across different metal ions, we compute the confusion matrix on all metal ions we studied. Predictors designated for one metal ion are tested on the other metal ions to compute the confusion matrix (Figure 11). To show the similarity between metal ions more clearly, we organized the order of the metal ions shown in the Figure 11 according to their physicochemical properties. Alkali metal ions (K^+^, Na^+^) and alkaline earth metal ions (Ca^2+^, Mg^2+^) are grouped together, while 3d transition metal ions (Mn^2+^, Fe^2+^, Fe^3+^, Co^2+^, Ni^2+^, Cu^2+^, Cu^+^, Zn^2+^) and d^10^ transition metals (Zn^2+^, Cd^2+^, Hg^2+^) are grouped together.

**Fig. 11.**
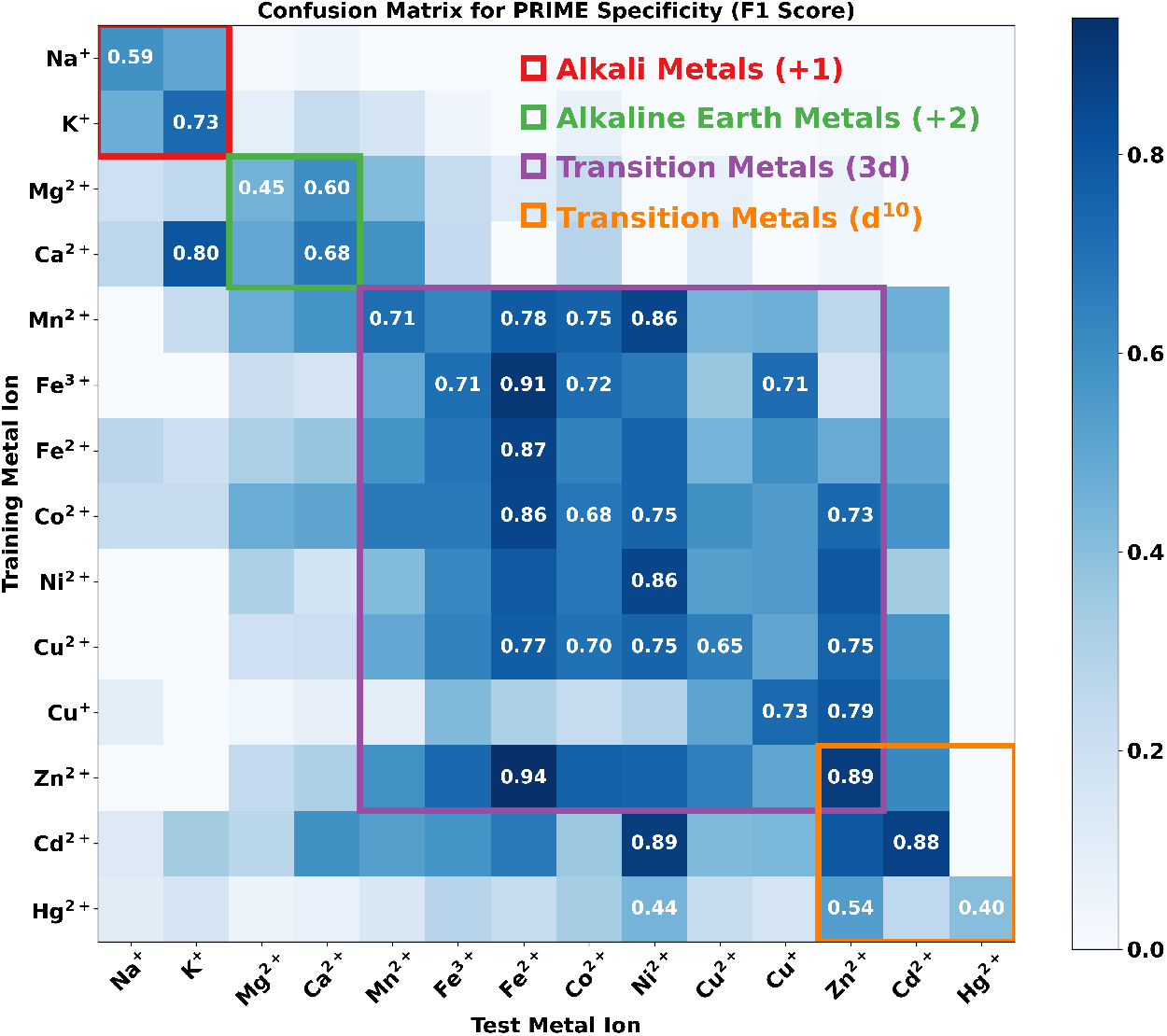
Heatmap of the confusion matrix of PRIME specificity across different metal ions. The Y-axis represents the predictors designated for a specific metal ion, while the X-axis represents the metal ion datasets used for testing. Values in the heatmap indicates the F1 scores of the prediction. To show the similarity between different metal ions more clearly, we organize the order of the metal ions according to their physicochemical properties. If a predictor achieves a higher F1 score on a metal ion that it was not trained on, the score is indicated in the corresponding cell.

Metal ions in the same group (e.g., alkali metal ions) show a clear compatibility with each other, i.e., the predictor trained on one metal ion (e.g., Na^+^) can achieve a high F1 score on the other metal ion (e.g., K^+^), indicating that the predictors take into account the chemicophysical properties of the metal ions. Some predictors, such as the Zn^2+^ predictor, show a very high F1 score on the prediction of Fe^2+^. This may be due to the similar binding patterns in proteins, as Zn^2+^ and Fe^2+^ both prefer to bind to histidine, glutamate, and aspartate residues (Supplementary Fig. S3), and have very similar van der Waals radii (0.74 Å for Zn^2+^ and 0.78 Å for Fe^2+^). It is very difficult to distinguish between these two metal ions, especially considering that the dataset for Fe^2+^ is not abundant compared to Zn^2+^ (153 vs. 441). In contrast, the predictor trained on the smaller Fe^2+^ dataset does not generalize well with respect to Zn^2+^ sites. This suggests that models trained on abundant data (such as for Zn^2+^) learn more general features that also apply to similar and less abundant ions. However, the reverse is not necessarily true.

### 2.8 Prediction speed of PRIME

We assessed the prediction speed of various methods, including BioMetAll, Metal3D, AllMetal3D, and PRIME. Sequence-based approaches were excluded from this comparison, as their speeds are generally comparable to PRIME-seq. The evaluation utilized a zinc-binding test set comprising 266 proteins, with an average of 13,250.72 ± 43,335.19 non-hetero atoms per protein. Predictions were restricted to zinc-binding sites, given the variability in supported metal ion types among methods. For instance, Metal3D predicts only zinc-binding sites, AllMetal3D supports predictions for 11 metal ions, and PRIME accommodates 14 metal ions. All evaluations were conducted on a Linux server equipped with 16 cores and an NVIDIA GeForce RTX 4090 GPU. While PRIME allows chain-specific predictions, this evaluation was performed at the whole-protein level to maintain consistency with other methods that lack chain-specific capabilities.

Structure-based methods generally require one to three minutes to predict a single protein, whereas PRIME completes this task in an average of 10.72 seconds (Figure 12A), demonstrating a substantial improvement in speed. This efficiency stems from two main factors: the effective integration of sequence and structural information, and a GPU-optimized probe generation algorithm. This avoids the exhaustive CPU-based search used by methods such as BioMetAll. The PRIME-probe module constitutes the primary contributor to inference time, operating approximately an order of magnitude slower than PRIME-seq due to the extensive processing of probes. Notably, probe predictions were augmented with five rotations, and inference time may vary depending on the number of rotations. The time required for probe generation and postprocessing is negligible and is therefore omitted from Figure 12B.

**Fig. 12.**
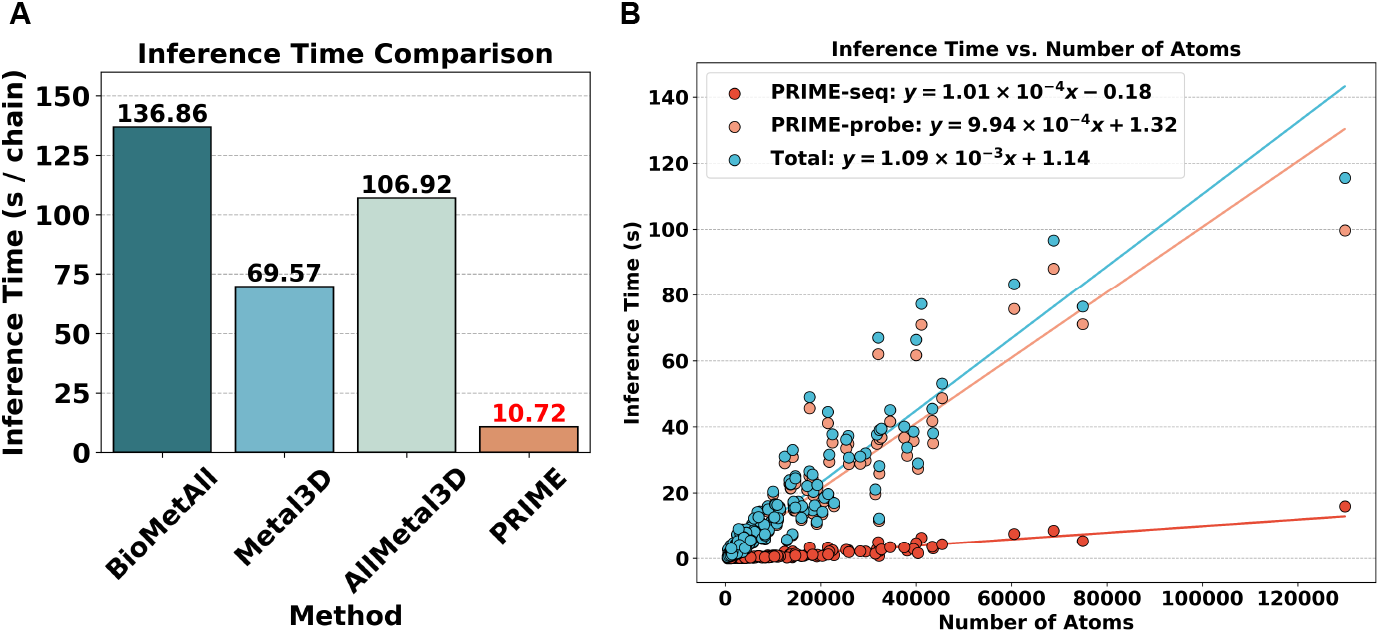
(A) The running time of different methods on the zinc-binding test set. The average running time per protein is indicated above the bars. (B) The relationships between inference time and the number of non-heteroatoms in the protein structures. Note that probe generation and postprocessing time are not included here as they are negligible.

## 3 Conclusion

We introduce PRIME, a novel deep learning framework for predicting metal-binding sites in proteins with high accuracy and efficiency. PRIME integrates both sequence and structural information, leveraging the power of representation learning to extract essential features from protein sequences using protein language models (PLMs) and from protein structures using pre-trained structure models (PSMs). The framework employs a probe generation algorithm to efficiently generate candidate binding sites based on geometric and evolutionary signals. PRIME achieves superior performance across a wide range of metal ions, including both abundant and less abundant types, and outperforms existing methods. Ablation analysis demonstrates that PSMs significantly enhance the accuracy of metal-binding site prediction. The prediction speed of PRIME surpasses that of existing structure-based methods, enabling its application in high-throughput metalloproteomics. These results underscore the potential of PRIME to advance the understanding of metalloproteins and their roles in biological processes, as well as their applications in biotechnology. While PRIME is currently trained and evaluated for single metal ions, future extensions will enable the prediction of binding sites accommodating multiple metal ions and metallocofactors.

## 4 Methods

### 4.1 Design of PRIME

PRIME is predicated on the rarity of metal-binding sites within protein structures. PRIME enhances both accuracy and efficiency by systematically excluding regions with low binding potentials. Reported structure-based methods employ physical constraints and statistical heuristics to address this issue [20], yet they remain neither sufficiently efficient nor accurate. We propose to adopt a neural network-based predictor (i.e., PRIME-seq) to discard the least apparent binding residues, followed by removal of geometrically impossible binding sites using structural information. Through a probe generation algorithm, we generate hundreds of candidate binding sites (probes). This amount is significantly less than the amount produced by BioMetAll, and achieve near-complete coverage of true binding sites.

Sequence-based methods typically predict the binding propensity of each residue, which indirectly indicates the ranking of likelihood of binding for each residue. Recent approaches, including LMetalSite [22] and M-Ionic [23], employ protein language models (PLMs) to extract high-dimensional representations of protein sequence, which are subsequently processed by additional neural network layers to predict residue-level propensities. Building upon similar concepts, we developed a sequence-based predictor, PRIME-seq, that performs comparably. While sequence-based methods lack structural awareness and fall short in prediction accuracy, we found them sufficient as rankers. We apply a very low threshold of ranking scores, recall of 99% on the validation set, to cover most binding residues. Residues surpassing this threshold are taken into account as candidate binding residues, which are then used in probe generation. This approach offers an effective balance between accuracy and efficiency of binding site prediction. When the number of predicted binding residues exceeding a certain threshold is insufficient, we select the minimum number of highest-ranked residues, determined by an emprical distribution based on chain length (Supplementary Fig. S9). In contrast to BioMetAll, we rely solely on distance-based geometric constraints to screen probes, a choice well-suited for parallel processing on modern graphical processing units (GPUs).

Structure-based methods struggle with the vastness of the search space for binding sites. PRIME addresses this challenge by combining sequence-based and structure-based strategies. While ESMBind [25] attempts to incorporate both types of information, integration was found to be weak, relying on separate handling and conventional energy minimization. SuperMetal [26] employs PLMs and structural input, yet was found to omit sequence features altogether; PLM representations are processed through a graph neural network (GNN) by SuperMetal, and candidate sites were generated via a diffusion process [51] that is both time-consuming and lacking in interpretability. In contrast, PRIME generates probes using sequence-derived features as well as geometric and statistical information, which are subsequently processed by another neural network (i.e., PRIME-probe) to predict the binding propensities of probes. This pipeline remains interpretable, tunable, and rich in intermediate outputs conducive to further analysis.

Structural pre-training has been widely employed to enhance performance across diverse downstream protein modeling tasks [52–56], yet it has not been applied to metal-binding site prediction. Structure-based masked language modeling (MLM), a standard pre-training objective, enables the network to learn geometric patterns inherent in protein structures, aiding metal-binding site prediction. In PRIME, we pretrain the PRIME-probe module on PDB data using MLM tasks across architecture ranging from 10 to 200 layers. Our findings demonstrate that such pretrained structure models (PSMs) substantially enhance prediction accuracy of metal-binding site prediction, providing new evidence for the value of pretraining in protein structural modeling.

### 4.2 Dataset

#### 4.2.1 Metal-binding data sets

We constructed a dataset based on the RCSB Protein Data Bank [2] (downloaded on 28th February 2025) and BioLiP2 [57] (released on 22nd February 2025). While the BioLiP2 dataset was curated with a rule of van der Waals radii between the metal ions and the coordinating atoms [57], we further verified these distances to ensure the quality of our dataset (Supplementary Fig. S10 and Table S4). To ensure the robustness of our method for wide-range metal-binding site prediction, we addressed the potential redundancy across datasets of different metal ions, which could compromise the integrity of training and validation by introducing information leakage from the test sets. Individual datasets were constructed for each metal ion, ensuring strict non-redundancy between training and test sets. For each metal ion, chains annotated by BioLiP2 were clustered at 30% sequence identity using MMseqs2 `easy-cluster` [58] with parameters `--min-seq-id 0.3 -c 0.8 --cov-mode 1`. Chains exceeding 2048 residues were excluded (Supplementary Fig. S11), thereby guaranteeing that no two structures share chains with more than 30% identity.

As the merged training and validation sets were also employed for hyperparameter tuning (Section 4.4), we further eliminated redundancy across metal ions. To preserve as many of the less abundant metal ions as possible, we began with the rarest metal ions we considered in this study (Supplementary Fig. S12) and incrementally incorporated more prevalent ions, excluding any chains redundant with the existing datasets. Details of this process are provided in Supplementary Algorithm S1. Following this procedure, we retained only metal ions with a sufficient number of annotated binding sites, namely, Zn^2+^, Ca^2+^, Mg^2+^, Mn^2+^, Fe^3+^, Cu^2+^, Fe^2+^, Co^2+^, Na^+^, Cu^+^, K^+^, Ni^2+^, Cd^2+^, and Hg^2+^. Summary statistics are presented in Supplementary Table S5. Other metal ions, though present in PDB structures, were omitted due to their extreme scarcity. For each metal ion, chains were randomly partitioned into training, validation, and test sets at an 8:1:1 ratio. The final dataset comprises 6,933 training chains, 865 validation chains, and 874 test chains (Supplementary Table S6). To ensure a fair comparison, redundant sequences were removed from our test set if they were found to overlap with the training sets of baseline methods. Redundancy was not removed against the test sets of AllMetal3D [40] as the complete dataset has not yet been released.

All protein structures were obtained from PDBj [59] in mmCIF format, and deprecated PDB IDs in BioLiP2 were retrieved automatically from the RCSB Protein Data Bank [2]. We retained only carbon, nitrogen, oxygen, and sulfur atoms within amino acid residues, removing all heteroatoms, including metal ions, water molecules, small molecule ligands, and nucleic acids. When multiple models were presented in the assembly, the first was selected for analysis.

### 4.2.2 Symmetrical mating

We constructed symmetrized protein structures using recorded symmetry mate data embedded with the crystal structure files. The PyMOL library `pymol`, specifically the `symexp` command, is employed to generate these symmetrized assemblies. For structure that fail to generate correctly or yield erroneous symmetrizations, we revert to the original mmCIF files. In certain cases, PyMOL erroneously fixes segments of the structure as part of the ensemble; to identify such instances, we evaluate the atomic density using the following expression:

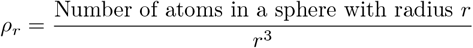

where *r* = 10 Å in this study. The constant 4*π/*3 is omitted from the denominator as its exclusion does not alter the efficacy of filtering when using an empirically determined threshold. We randomly sample ten spherical regions with a 10-Å radius within each protein, with replacement, and compute the average density 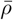. Structures exhibiting 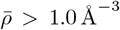 are excluded, and their corresponding raw mmCIF structures are utilized instead. The resulting symmetrized structures are incorporated during training, validation, and testing.

#### 4.2.3 Pretraining data sets

Only X-ray crystal structures with high resolution (≤2.0 Å) available in RCSB protein data bank are employed for pretraining. These structures are clustered using the same MMseqs2 settings described above, ensuring that no chains in the training and validation sets share more than 30% sequence identity. From the resulting non-redundant dataset, 1,000 structures were randomly selected for validation, while the remaining 16,840 were reserved for training. The test was omitted, as the pretraining phase is not intended for comparative evaluation against other methods.

### 4.3 PRIME

#### 4.3.1 PRIME-seq

PRIME-seq, a sequence-based predictor within the PRIME framework, leverages protein language models (PLMs) to encode sequences with both efficiency and precision. Larger PLMs typically yield richer representations imbued with greater generalization capacity than their smaller counterparts; however, the elevated dimensionality of such representations can increase the risk of overfitting in downstream applications with limited fine-tuning data. The selection of an appropriate PLM is thus contingent upon the specific task and dataset [37, 39]. We employ the ESM2 series and ProtTrans models to extract high-dimensional representations of protein sequences, selecting the three largest ESM variants (650M, 3B, 15B parameters) along with ProtXL-UR50 for hyperparameter tuning. These representations are subsequently passed through an interaction layer designed to capture residue-residue relationships, followed by a fully connected (FC) layer that predicts the binding propensity of each residue.

Various configurations involving PLMs, RNNs including a bidirectional gated recurrent unit (GRU) [60], long short-term memory (LSTM) [61] networks, and transformer encoder layers [62] with hidden sizes of [128, 256, 512] and layer numbers of [2, 3, 4] were examined. The AdamW optimizer [63] was employed with a learning rate of 0.001 and a batch size of 32. A cosine annealing schedule was used to adjust the learning rate. Training was terminated early if the validation AUC failed to increase for five consecutive epochs or if it reached 100 epochs.

A shared model for all metal ions was initially trained using the merged training and validation sets to capture the general metal-binding propensity of protein sequences. After training the shared model, it was fine-tuned on the training set of each metal ion. PRIME-seq employs an additional fully connected (FC) layer appended to the interaction layer to predict the binding propensity *p*_*m*_, where *m* denotes the metal ion. The output from the shared FC layer, *p*_shared_, is also incorporated, and the final binding propensity was computed as *p* = *p*_*m*_ + *p*_shared_. During shared model training, the metal-specific FC layer is not used, whereas in the fine-tuning stage, all parameters, including the newly added FC layer, are trainable.

This design integrates data across all metal ions, necessitating meticulous measures to prevent data leakage between ions. For instance, the test set of LMetalSite includes a Mg^2+^ binding site from protein 7CEN A, which shares a highly homologous Mn^2+^ binding site with 5VNU A in its training set (RMSD: 0.281 Å). To mitigate such issues, separate train/test sets for each metal ion are avoided. Instead, a gradual paradigm is employed to construct the training, validation, and test sets, ensuring non-redundancy across ions (Supplementary Algorithm S1).

#### 4.3.2 Ranking binding residues

The sequence predictor assigns each residue in a protein a binding propensity score between 0 and 1. This score serves as a ranking metric for binding residues, aligning with the definition of AUC (Area Under the Curve), which corresponds to the probability that a randomly selected positive residue exhibits a higher score than a randomly selected negative one. Existing methods, including ours, typically optimize the AUC as a principal performance indicator. All residues in the sequence are ranked by their predicted propensity to bind metal ions and the top *N* residues are selected for subsequent structure-based analysis. The value of *N* is computed as

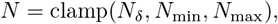

where *N*_*δ*_ denotes the number of residues with predicted propensity ≥ *δ*, and *δ* represents the highest threshold that preserves recall ≥ 0.99 on the validation set (Supplementary Table S7). The lower bound *N*_min_ is a function *f* that increases with the length of the protein sequence, while the upper bound *N*_max_ is a constant set to 400 by default. We adopt the form *f*_*a,λ,b*_(*x*) = *ax*^*λ*^ + *b*, where *x* is the sequence length, and the goal is to approximate the number of binding residues in a single protein chain. To optimize the parameters *a, λ*, and *b*, we minimize the following loss function using the `scipy.optimize.minimize` method with the `L-BFGS-B` algorithm [64]:

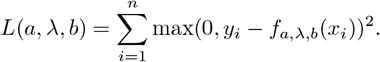

Here, *n* is the total number of chains, *y*_*i*_ is the number of binding residues in the *i*-th chain, and *x*_*i*_ is its sequence length. Optimization is conducted with initial values of *a* = 1, *λ* = 0.5, and *b* = 0, constrained within (0, ∞), (0, 1), and (∞, ∞), respectively. Given the pronounced long-tail distribution in binding residue counts (Supplementary Fig. S9), only the short half of the chains in the training and validation sets are used. The fitted parameters are *a* = 1.8218, *λ* = 0.6250, and *b* = 0.0629.

#### 4.3.3 Probe generation algorithm

We introduce a probe generation algorithm designed to identify candidate binding sites within protein structures. Binding ions are frequently coordinated by nitrogen, oxygen, and sulfur atoms in amino acid residues, particularly in the case of transition metals. The algorithm initiates by filtering positions containing such atoms. For each metal, we consider the the amino acids whose participation in metal-binding exceeds 0.1% among all binding residues (Supplementary Fig. S3). The N/O/S atoms within these residues are collected, and gridpoint boxes with 5 Å margins and 0.5 Å spacing are generated around each residue, encompassing the relevant atoms. These gridpoints, referred to as “probes”, are then filtered by the following criteria:

- Each probe must maintain a minimum distance *δ*_min_ from any protein atom (including C/N/O/S), expressed as:

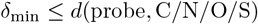
- Probes must lie within a specific range from N/O/S atoms, expressed as:

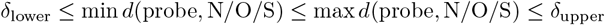

The thresholds *δ*_min_, *δ*_lower_, and *δ*_upper_ are empirically derived from the validation set. We calculate the distance distribution of N/O/S atoms to bound metal ions (Supplementary Fig. S13), which reveals a pronounced clustering. Accordingly, we determine the 25th and 75th percentiles, denoted as *δ*_0.25_ and *δ*_0.75_. The interquartile range (IQR) is then defined as IQR = max(*δ*_0.75_ −*δ*_0.25_, 0.5), where minimum IQR 0.5 is enforced to compensate for sparse data. The distance bounds are set as:

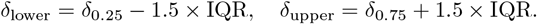

Note that *δ*_lower_ and *δ*_upper_ are specific to both metal ions and N/O/S atomic types. We define *δ*_min_(*m*) = min_*a*_ *δ*_lower_(*a, m*) for each metal ion *m*, where *a* ranges over all N/O/S atoms.

In the subsequent step, probes are clustered to reduce redundancy. A probe is considered to be “touching” an atom if its distances to that atom lies within the interval *δ*_lower_ to *δ*_upper_. Probes that touch exactly the same set of N/O/S atoms are grouped into a cluster *C*. For each cluster *C* ⊂ ℝ^3^ of probes, we compute the geometric center to serve as its representative. Each probe carries two additional attributes: the maximum predicted binding propensity *kp* of the residue it touches and highest portion *ks* among the N/O/S atoms it touches. These attributes are inherited by the representative probe. Representative probes are ranked in descending order of *ks*. Probes within a distance less than *d*_max offset_*/*2 of higher-ranking probes are discarded (*d*_max offset_ = 3.0 Å for Zn^2+^ and 5.0 Å for others). The full pseudocode of the algorithm is provided in Supplementary Algorithm S2 and S3. The final set of probes, processed through these algorithms, serves as the input for PRIME-probe.

In PyTorch, Boolean tensors are stored with 1 byte (8 bits) per element due to technical constraints, which can lead to inefficiencies for large tensors. To optimize memory usage, these tensors are packed into more compact byte tensors, achieving approximately 8x memory savings and enabling more efficient computation. The packing and unpacking processes are implemented using vectorized bitwise operations, as detailed in Supplementary Algorithms S4 and S5.

#### 4.3.4 PRIME-probe

##### Pretraining of structure predictor

We adopt a masked language modeling (MLM) task to pretrain the structure predictor. A single alpha carbon atom within a protein backbone is randomly selected, around which a cubic region measuring 10 Å per side is defined. All carbon, nitrogen, oxygen, and sulfur atoms within this cube are excised from the structure. The entire residue containing the selected alpha carbon is masked, and the remaining atoms are provided as input to the structure predictor, which outputs a 20-dimensional vector. The MLM objective is to infer the amino acid type of the masked residue. Model training is conducted using cross-entropy loss and the AdamW optimizer.

The architecture of the structure predictor is a three-dimensional residual network (ResNet3D) with depths of 10, 18, 34, 50, 101, 152, or 200 layers, mirroring the canonical ResNet design [41] while being adapted to accommodate volumetric data (Figure 13A-E). Input features comprise atomic coordinates and element types (C, N, O, or S). A 60 ×60 ×60 grid is constructed around the alpha carbon with a spacing of 0.3 Å in each dimension. Grid features are computed by applying a Gaussian kernel with standard deviation *σ* = 0.18, resulting in blurred representations of shape [batch size, 60, 60, 60, 4], where 60 denotes the number of grid points along each axis, and 4 corresponds to the number of atom types. These features are treated as 3D images and passed into the ResNet3D. Pretraining is performed with a batch size of 32, a weight decay of 0.01, and a cosine annealing schedule over 1,000 epochs. The learning rate is linearly warmed up from 10^−5^ to 10^−3^ during the initial 10% of training, and subsequently decayed to 10^−5^ over the remaining epochs. Data augmentation via coordinate rotation is omitted, as the long duration of pretraining allows the model to capture the geometric patterns across orientations. Experiments across different ResNet3D sizes revealed improved accuracy with increasing depth (Supplementary Fig. S6). These validated accuracies are comparable to those reported in prior studies on protein masked language modeling [52–54, 56].

**Fig. 13.**
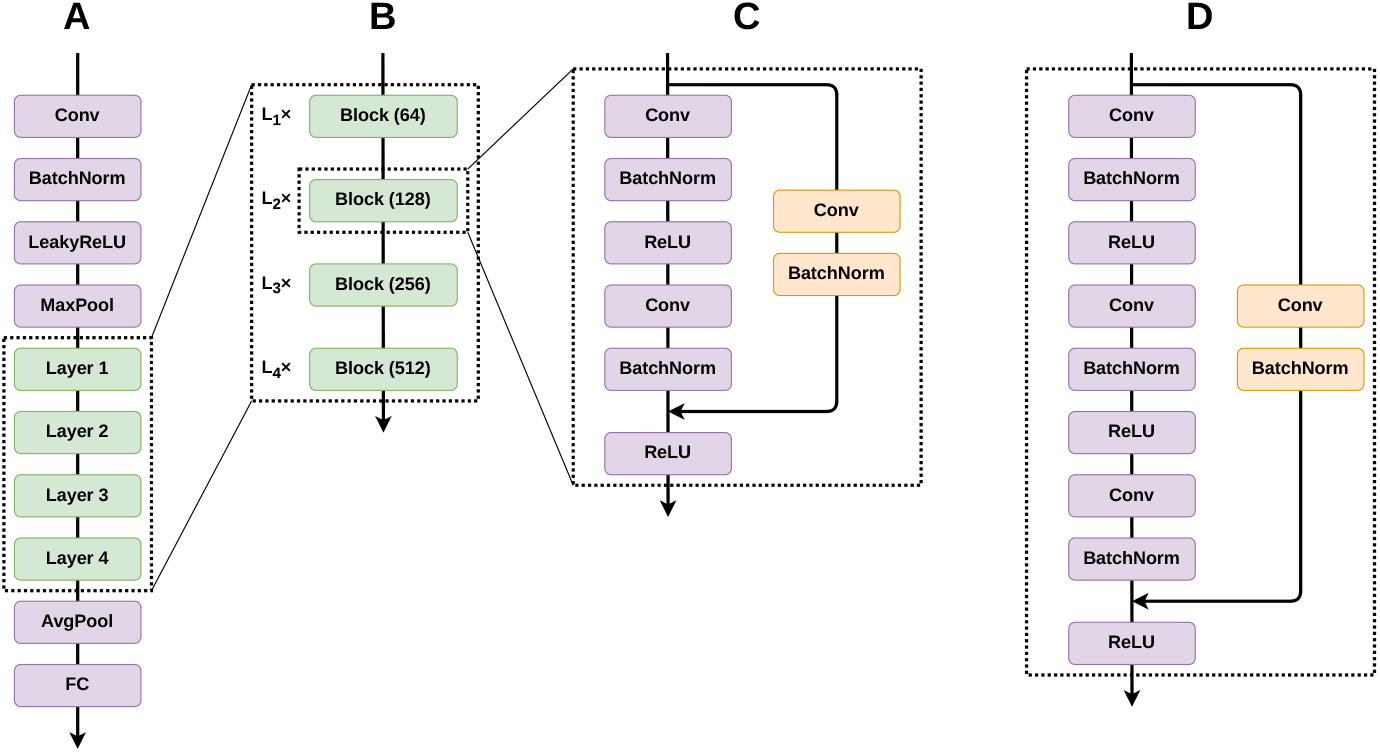
(A) ResNet3D consists of multiple stacked convolutional layers (Conv), batch normalization layers (BatchNorm), activation functions (LeakyReLU and ReLU), pooling layers (MaxPool and AvgPool), and fully connected layers (FC). Four residual blocks are included, each with two distinct types: (B) Layers 1–4 comprise stacked blocks with channel sizes ranging from 64 to 512. The blocks are of two types: (C) the Basic Block, used in shallower networks (ResNet3D-10, -18, -34, -50), and (D) the Bottleneck, designed for deeper networks (ResNet3D-101, 152, 200). Detailed parameters of ResNet3D are listed in Supplementary Table S8.

##### Fine-tuning on metal-binding prediction

The pretrained weights of ResNet3D are used to initialize the structure predictor, with the exception of the output linear layers, which differ in dimensionality between pretraining (20 output dimensions) and fine-tuning (2 output dimensions) and are thus randomly initialized using uniform distributions. A learning rate of 10^−4^ is employed for fine-tuning the pretrained model. For baseline comparisons with randomly initialized ResNet3D, a learning rate of 10^−3^, which is the same as the value used in pretraining, is applied. The model is fine-tuned using the same batch size and optimization settings as in pretraining, trained for 100 epochs with an early stopping patience of 5 for abundant metal ions (Zn^2+^, Ca^2+^, Mg^2+^, Mn^2+^, Fe^3+^, Cu^2+^), and for 200 epochs with a patience of 20 for the remaining, less abundant metal ions. Data augmentation is introduced during fine-tuning by applying random rotations to the atomic coordinates in each training step. Rotation matrices are constructed using uniformly sampled angles along the *x, y*, and *z* axes in the range [0, 2*π*].

The training objective of the structure predictor comprises two components: a cross-entropy loss assessing whether a probe is a metal-binding site for a give metal ions, and a smooth *L*_1_ loss measuring the distance between the probe and the metal ion in positive samples. The cross-entropy loss will be discussed in section 4.3.5. The smooth *L*_1_ loss [65] offers stable gradient behavior and robustness to outliers, making it popular and widely used in object detection literature [65, 66]:

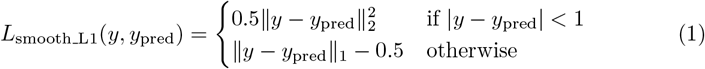

#### 4.3.5 Hard mining and reweighting strategy

To enhance the performance of the structure predictor, we adopt a hard mining and reweighting strategy. Originally introduced in the field of object detection [67], hard mining emphasizes challenging samples during training, leading to improved prediction performance. In each training step, negative probes with the highest predicted probabilities are selected as “hard samples”, while the reaminder are discarded. Next, [67], we retain the top 20% and fine-tune the structure predictor on zinc-binding proteins to assess its efficacy. As illustrated in Supplementary Table S9 and Fig. S14, the best F1 score on the validation set is achieved with the hard mining strategy, underscoring its effectiveness as a structure predictor.

We use a reweighting strategy to address the severe class imbalance between positive and negative probes. Standard cross-entropy loss performs poorly in such scenarios because it implicitly assumes balanced classes. Therefore, we adjust the loss function to give more weight to the under-represented positive probes. To reduce potential bias, we compute the total loss by separating positive and negative contributions: *L*_total_ = *L*_positive_+*αL*_negative_, where *α* denotes the weighting factor for negative probes. In this study, we fix *α* = 5 across all metal ions. In addition, as the distance loss is essentially the MAE (mean absolute error) or RMSE (root mean square error) of the distance between the probe and the metal ion, ranging over several angstroms during training and thus substantially larger than the cross-entropy loss, we also reweight the smooth *L*_1_ loss by a factor of 0.1. The final loss function is:

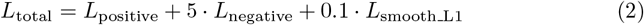

#### 4.3.6 Postprocessing

For the final prediction, a postprocessing procedure was applied following the training of the probe predictors. To enhance robustness, each probe was augmented by rotating it around its center with random angles uniformly sampled from [0, 2*π*] in each dimension, repeated a fixed number of times (default: 5). The binding probability for each probe was then determined as the maximum across its augmented counterparts. Subsequently, `scipy.cluster.hierarchy.fclusterdata` was employed to cluster the probes within a defined threshold (3.0 Å for all metals, unless otherwise specified), using the `criterion=’distance’` option. Clusters were retained if they contained multiple probes (≥2) and any predicted probability in the cluster exceeded the threshold (default: 0.5). Within each retained cluster, the probe with the highest probability was selected as the final prediction (Supplementary Algorithm S6).

### 4.4 Hyperparameter Tuning

The distribution of validated and test AUC values achieved by PLMs is presented in Supplementary Fig. S15. The entire dataset encompassing all metal ions was utilized for hyperparameter optimization. While all PLMs demonstrate comparable validated AUCs, their test AUC distributions exhibit notable differences. Interestingly, the smallest PLM, ESM T33, achieves the second-highest validated AUC. Given the computational efficiency of smaller PLMs, ESM T33 was selected for PRIME-seq. Supplementary Fig. S16 illustrates the performance of various interaction layer types, with the gated recurrent unit (GRU) layer significantly outperforming others on the validation set. On the test set, GRU also achieves the highest AUC, except in the case of ESM T36. These results indicate that GRU, as a relatively lightweight neural network, performs robustly, whereas more complex architectures such as LSTM and transformer layers may require larger datasets to fully exploit their potential. The influence of hidden sizes and layer numbers is depicted in Supplementary Fig. S17. Although GRU consistently outperforms other interaction layers, variations in hidden sizes and layer numbers do not lead to significant performance differences. This study employs a GRU interaction layer with a hidden size of 256 and three layers (Supplementary Table S10), as it achieves the highest AUC on the validation set while maintaining a reasonable model size.

For the structure predictor, the pretrained ResNet3D models were fine-tuned on the training and validation sets of zinc-binding proteins to identify the optimal depth (Supplementary Table S9). The hyperparameters, including ResNet3D layers, pre-training strategies, and loss functions, were systematically optimized. Notably, the pretrained ResNet3D models were found to consistently outperform those trained from scratch, highlighting the efficacy of pretraining in enhancing model performance. Among the configurations, the ResNet3D-152 model trained with the hard mining strategy achieved the highest validated F1 score and was therefore selected for PRIME-probe. Due to computational constraints, further evaluations on other metal ions were not conducted, and the ResNet3D-152 model was uniformly applied across all metal ions.

### 4.5 Comparison with other methods

PRIME comprises two predictors, PRIME-seq and PRIME-probe, each benchmarked against leading sequence-based and structure-based approaches, respectively. PRIME-seq is assessed through a residue-level metal-binding prediction task, to determine whether a given residue participates in metal coordination. On the other hand, PRIME-probe is evaluated at the structural level, where both the positions and types of metal-binding sites are predicted.

PRIME-seq is compared with state-of-the-art sequence-based methods, including LMetalSite [22], MetalNet [68], and M-Ionic [23]. For structure-based comparisons, PRIME-probe is evaluated against Metal3D [12] on Zn^2+^ and AllMetal3D [40] for more metal ions, as well as BioMetAll [20]. ESMBind [25] is excluded from comparison, as its assumption of a single binding site diverges from our task. Similarly, AlphaFill [29] requires prior knowledge of binding site presence, making it unsuitable for evaluation. SuperMetal [26] and MetalSiteHunter [32] are not evaluated because they lack a complete prediction pipeline. MIB [18] lacks both standalone software and an accessible web service. MIB2 [19] only offers an online service with strict query limitations, rendering it impractical for evaluating over 800 proteins in our test set. The performance of these systems was previously reported to be inferior to Metal3D, and thus we omitted a comparison with them. Mebipred [21], which predicts metal-binding propensity at the protein level, addresses a different task and is therefore excluded from our comparison. For locally executable structure-based methods, a maximum runtime of ten-minutes per protein is enforced, within which performance metrics are recorded. The time required for MSA (multiple sequence alignment) generation by MetalNet is not included in this limit.

For LMetalSite, we utilized the official GitHub repository https://github.com/biomed-AI/LMetalSite. MetalNet was obtained from https://github.com/wangchulab/MetalNet, with multiple sequence alignments constructed using `hhblits` from HH-suite3 [69] and the Uniclust30 [70] (version: 2023_02), excuted with the command `hhblits -i <query.fasta> -d <Uniclust30 database> -oa3m <output.a3m> -n 3 -cpu 32`. M-Ionic was obtained via its repository https://github.com/TeamSundar/m-ionic.Metal3D was obtained from https://github.com/lcbc-epfl/metal-site-prediction while AllMetal3D and BioMetAll were installed through the Python packages `allmetal3d` and `biometall`, respectively. BioMetAll retrieves structures in PDB format from the RCSB Protein Data Bank or converts mmCIF files to PDB using the Bio.PDB module if remote retrieval fails.

### 4.6 Evaluation metrics

Metal-binding site prediction differs from standard binary classification tasks, as true negatives (TN) cannot be defined without a clear “non-binding” state in protein structures. True positives (TP) are predictions within 5.0 Å of an actual metal-binding site, following established methodology [12]. Classification performance is evaluated using true positives (TP), false positives (FP), and false negatives (FN), from which additional metrics are derived. The F1 score is defined as

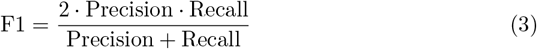

where precision is

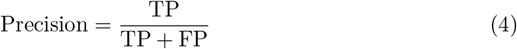

and recall is

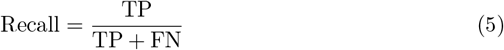

AUC (Area Under the Receiver Operating Characteristic Curve) was also used to evaluate the performance of PRIME-seq, calculated using the algorithm from the `sklearn.metrics` package.

To evaluate the performance of the structure predictor in the position prediction of binding sites, we additionally compute the root mean square error (RMSE), defined as:

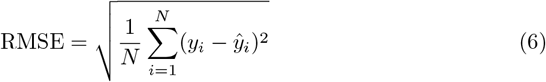

where *y*_*i*_ and *ŷ*_*i*_ denote the actual and predicted metal positions, respectively.

## 5 Data availability

All dataset used to train and evaluate the models and the data required to reproduce figures and tables in this manuscript have been deposited in Zenodo under https://doi.org/xxx.

## 6 Code Availability

All datasets and corresponding source code utilized in this study are publicly accessible at https://github.com/xu-shi-jie/prime. Additionally, a web server for PRIME is hosted at https://onodalab.ees.hokudai.ac.jp/prime.

## 7 Author Contributions

All authors conceived and designed the study. S.X. developed the method, performed data analysis, and wrote the paper. A.O. supervised the research project and critically reviewed and revised the manuscript. All authors approved the final manuscript.

## Acknowledgements

This work was supported by Hokkaido University DX Doctoral Fellowship (JST SPRING, Grant Number JPMJSP2119) to S.X. and JSPS KAKENHI Grant Number JP24H02213 in Transformative Research Areas (A) JP24A202 Integrated Science of Synthesis by Chemical Structure Reprogramming (SReP) and JP24K01533, and SATREPS Project “Recovering High-Value Bioproducts for Sustainable Fisheries in Chile (ReBiS)” funded by JST/JICA (Grant Number JPMJSA2206) to A.O.

