## Supplementary information for "Probe-Based Identification of Metal-Binding Sites Using Deep Learning Representations"

### Supplementary Material

Shijie Xu[1,2] and Akira Onoda [1,2,*]

[1]Graduate School of Environmental Science, Hokkaido University, Sapporo, Hokkaido, Japan
[2]Faculty of Environmental Earth Science, Hokkaido University, Sapporo, Hokkaido, Japan

*Corresponding author.

October 4, 2025

### 1 Supplementary figures

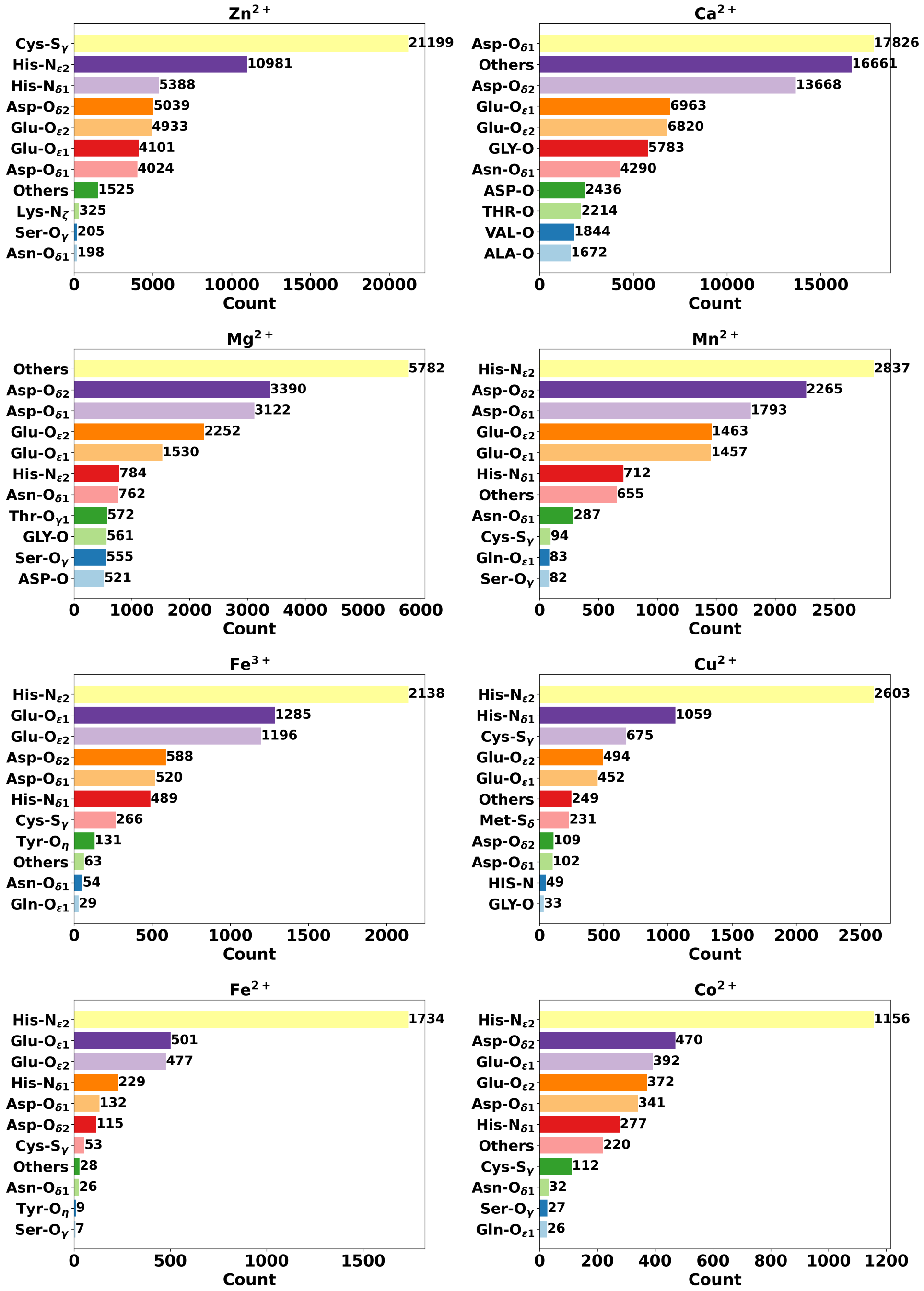


**Fig. S1** The distribution of atom-metal binding pairs in the BioLiP2 dataset. Presented metal ions include $Zn^{2+}$, $Ca^{2+}$, $Mg^{2+}$, $Mn^{2+}$, $Fe^{3+}$, $Cu^{2+}$, $Fe^{2+}$, and $Co^{2+}$.

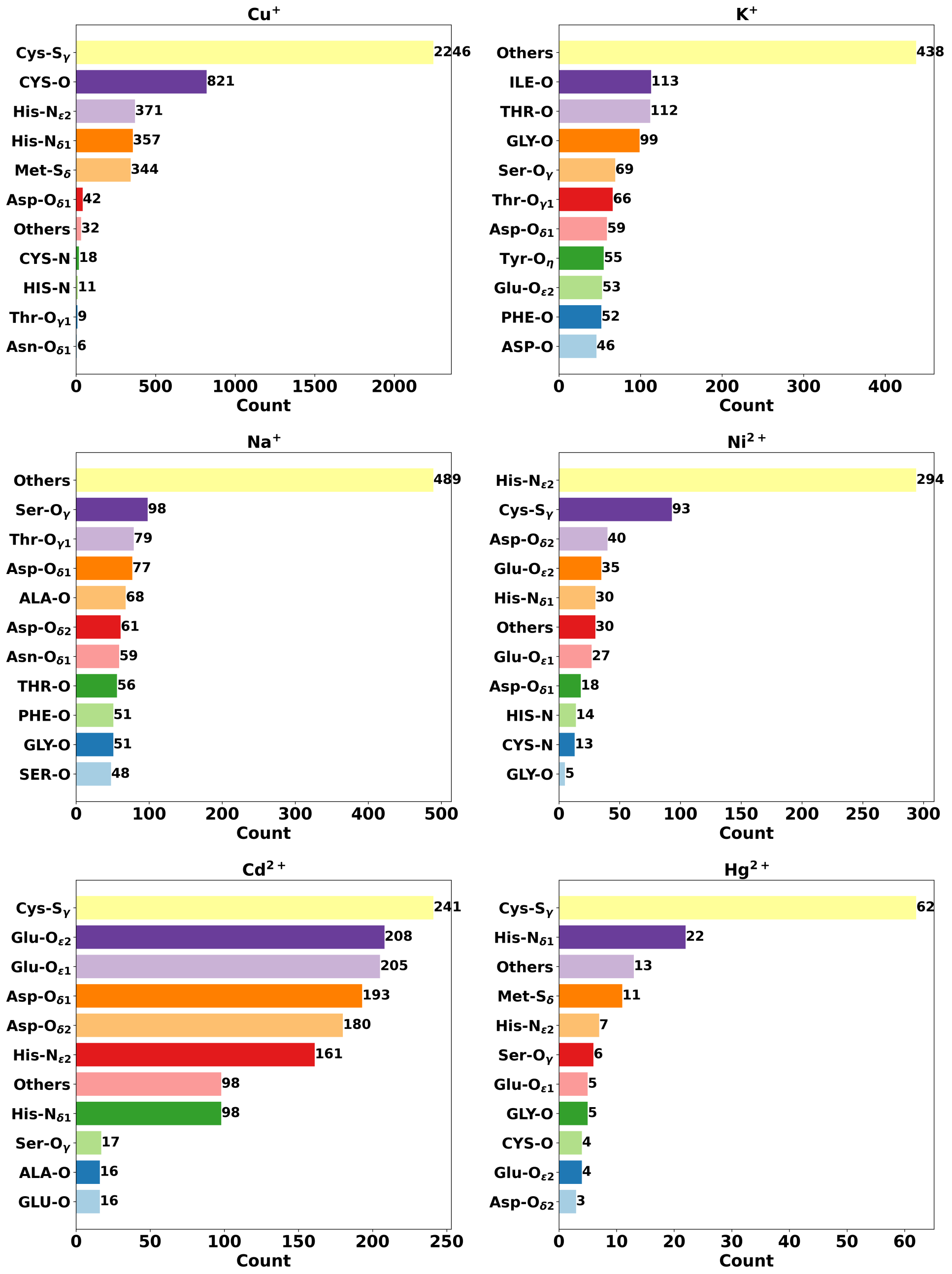


**Fig. S2** The distribution of atom-metal binding pairs in the BioLiP2 dataset. The remaining metals, including $Cu^{+}$, $K^{+}$, $Na^{+}$, $Ni^{2+}$, $Cd^{2+}$, and $Hg^{2+}$, are presented.

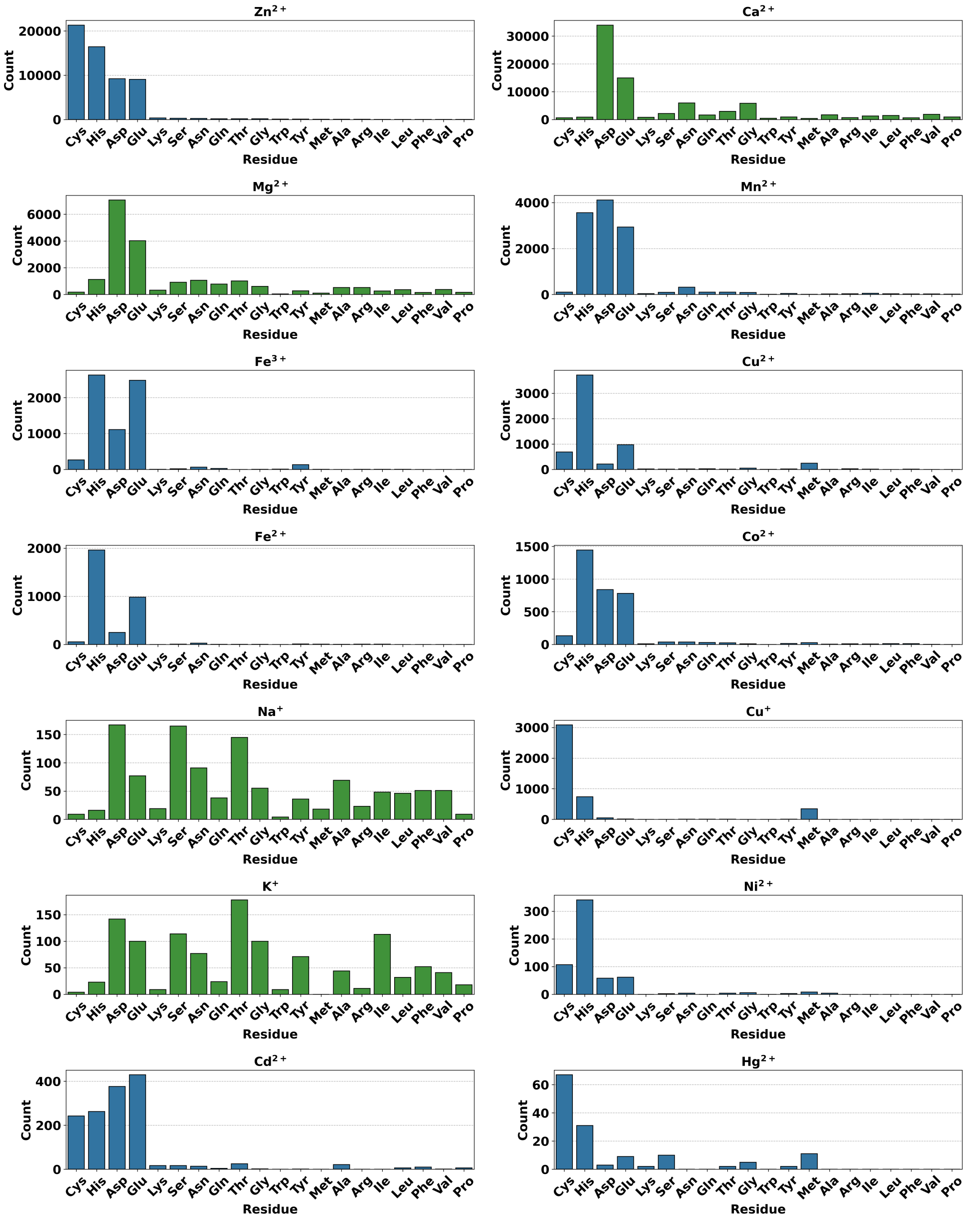


**Fig. S3** The distribution of residue-metal binding pairs in the BioLiP2 dataset. Each subfigure corresponds to a specific metal ion. Blue bars denote transition metals, while green bars represent alkali and alkaline earth metals. The x-axis specifies residue types, and the y-axis indicates the count of binding residues. Residues are ordered by their binding counts on the zinc binding site.

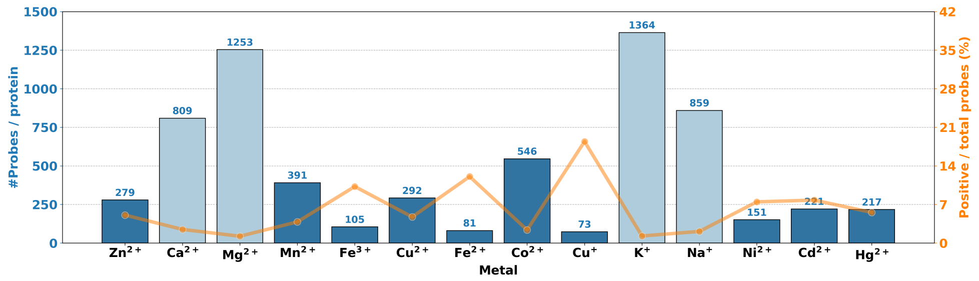


**Fig. S4** The number of probes per protein and the proportion of positive probes. The left y-axis displays the number of probes per protein, with dark blue bars representing transition metals and light blue bars denoting alkali and alkaline earth metals. The right y-axis, indicated by the orange curve, illustrates the proportion of positive probes among all probes.

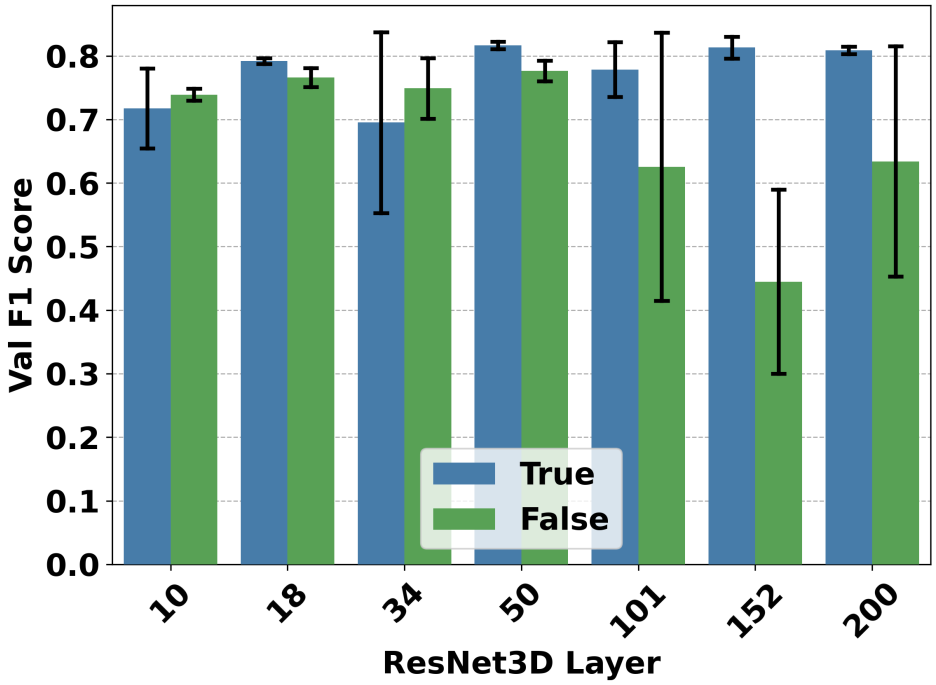


**Fig. S5** Performance of PRIME on zinc-binding prediction across varying ResNet3D layers and pretraining strategies. The y-axis indicates the validated F1 score, while the x-axis denotes the number of ResNet3D layers. Blue bars represent models with pretraining, and green bars correspond to models without pretraining.

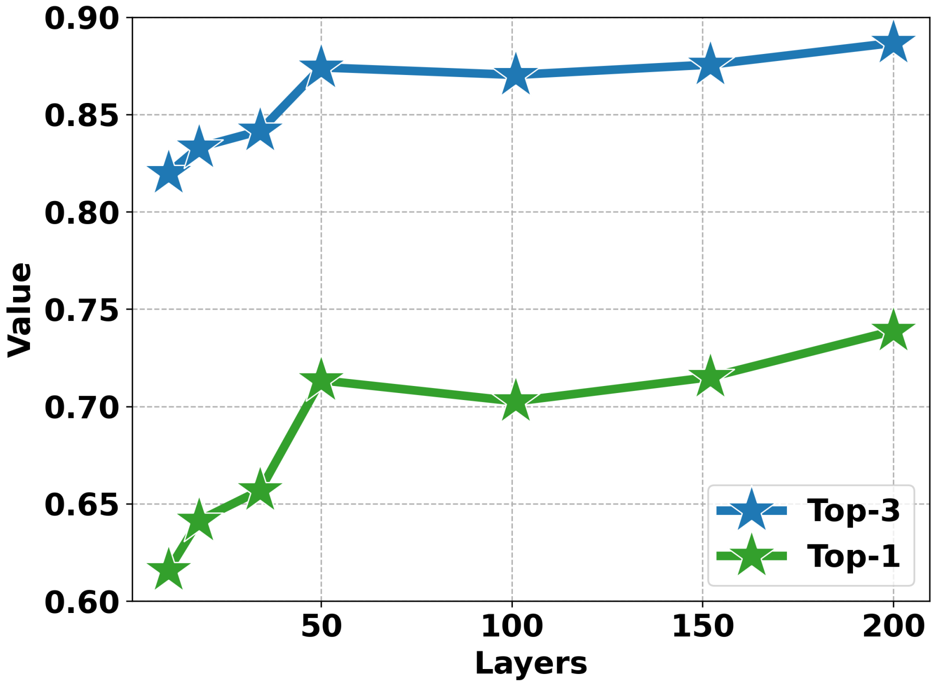


**Fig. S6** The top-1 and top-3 accuracies of the masked language modeling task for ResNet3D with varying depths: 10, 18, 34, 50, 101, 152, and 200 layers. The x-axis denotes the number of ResNet3D layers, while the y-axis represents the accuracy. Top-1 accuracy is depicted in green, and top-3 accuracy in blue. The curves reveal an inflection point around 50 layers, potentially deviating from the scaling law due to the constrained size of the pretraining dataset.

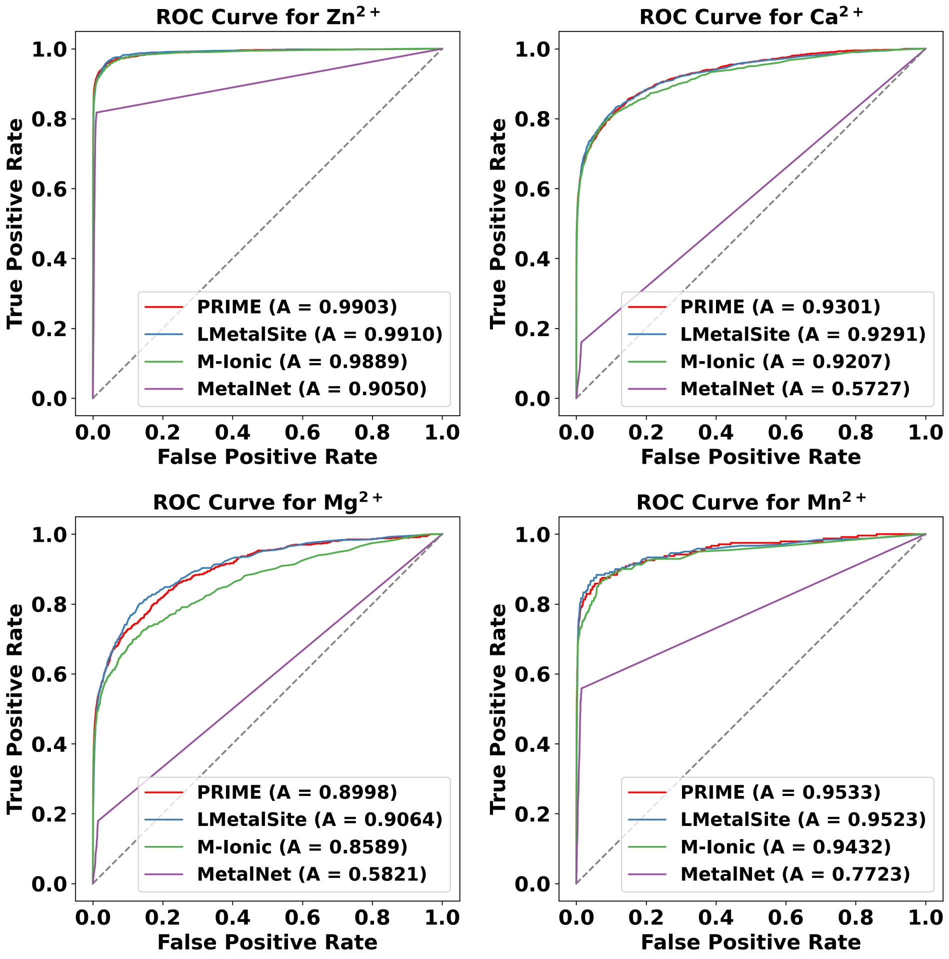


**Fig. S7** The ROC curves for the prediction of metal-binding residues by PRIME and other sequence-based methods for $Zn^{2+}$, $Ca^{2+}$, $Mg^{2+}$, and $Mn^{2+}$. The legends indicate the AUC values of each method.

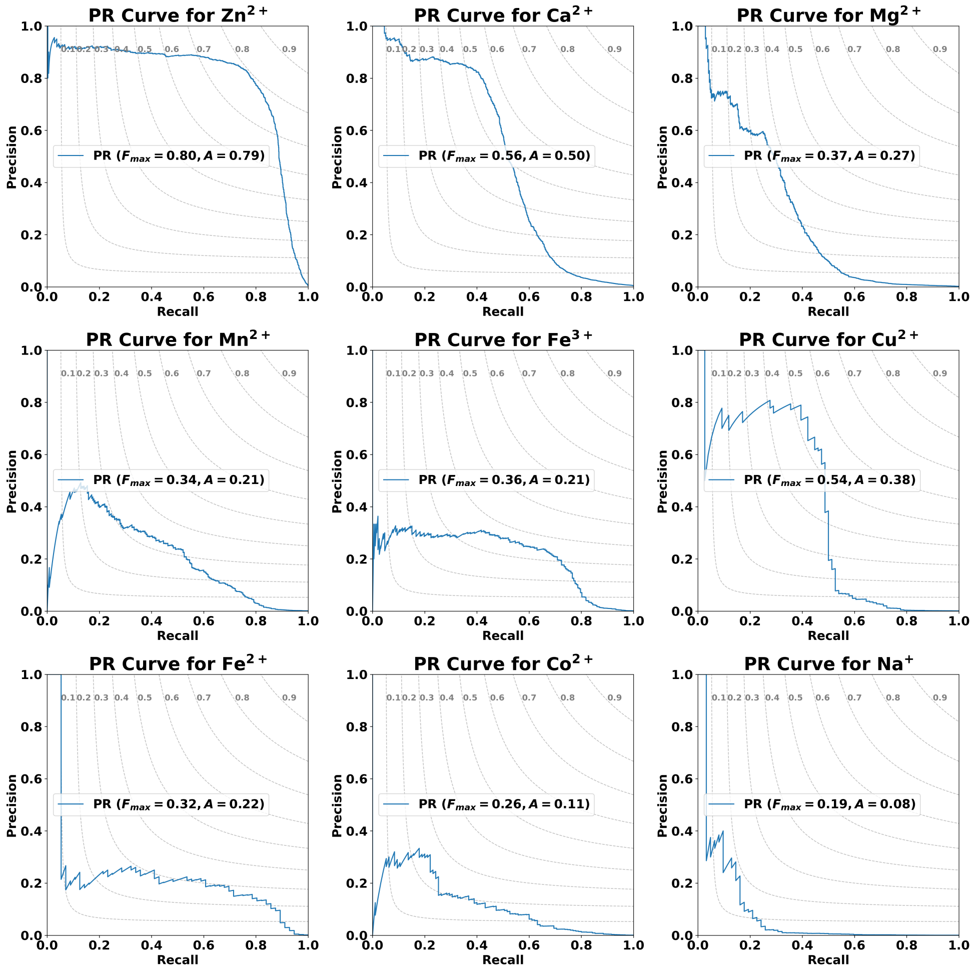


**Fig. S8** The precision-recall curve of PRIME-seq on the test binding sets for $Zn^{2+}$, $Ca^{2+}$, $Mg^{2+}$, $Mn^{2+}$, $Fe^{3+}$, $Cu^{2+}$, $Fe^{2+}$, $Co^{2+}$, $Cu^{+}$, $K^{+}$, and $Na^{+}$. Other metals, including $Ni^{2+}$, $Cd^{2+}$, and $Hg^{2+}$, are excluded due to their curves being nearly indistinguishable from the x-axis.

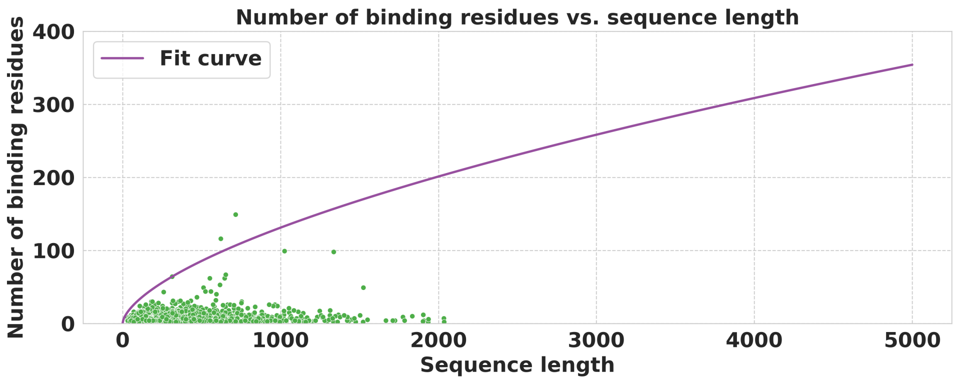


**Fig. S9** The relationship between the number of binding residues in a protein and its sequence length in the validation set. The y-axis indicates the number of residues involved in metal coordination, irrespective of the metal type, while the x-axis represents the sequence length of the chain. Green dots denote all chains with binding residues, and the purple curve illustrates the fitted function $y = a \cdot x^{\lambda} + b$, which minimally encompasses the first half of the data points. Further details are provided in the main text.

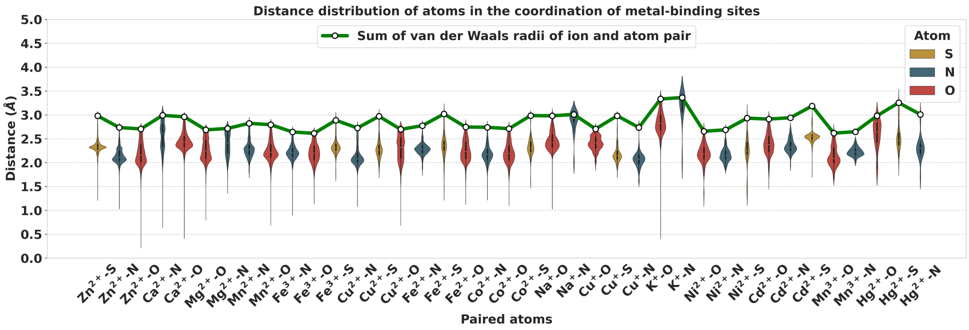


**Fig. S10** The distribution of distances between binding atoms and metal ions. The y-axis denotes the distance, while the x-axis categorizes ion-atom pairs, with sulfur atoms in yellow, nitrogen in blue, and oxygen in red. Green curves represent the sum of van der Waals radii for metal ions and atoms, delineating a clear upper bound for the distance distribution.

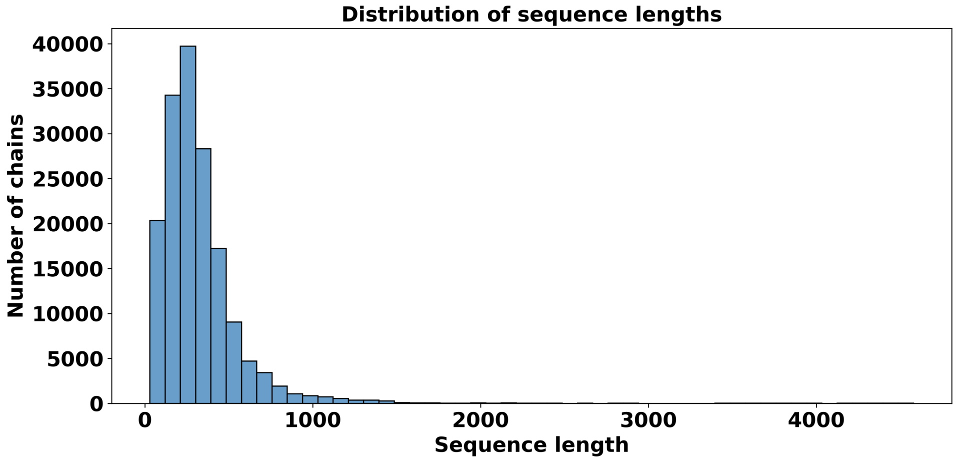


**Fig. S11** The distribution of chain lengths in the BioLiP2 dataset. Most chains contain fewer than 2,000 residues, a threshold adopted in this study for selecting chains for training purposes.

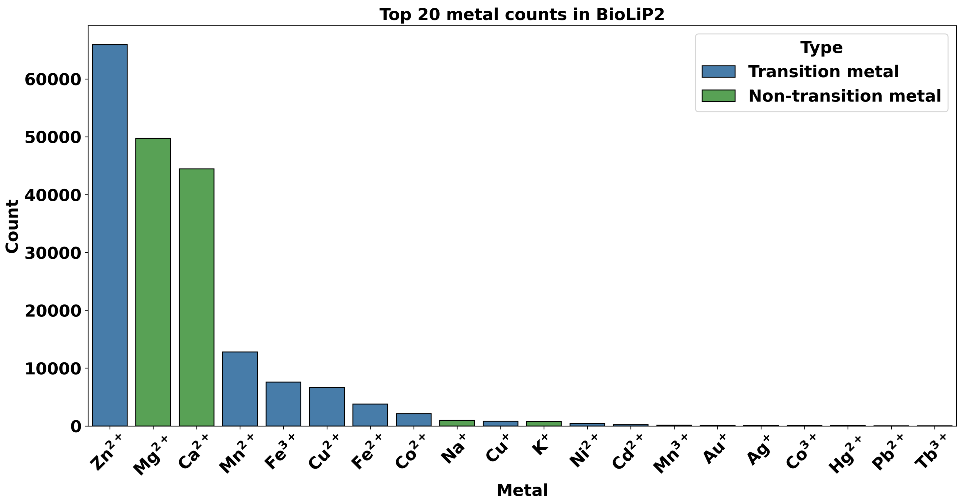


**Fig. S12** The distribution of the top 20 most abundant metal ions in the BioLiP2 dataset. Blue bars represent transition metals, while green bars denote alkali and alkaline earth metals. We selected the top 13 most abundant metal ions, along with $Hg^{2+}$, for training and evaluation, as the remaining ions are insufficiently represented for robust training.

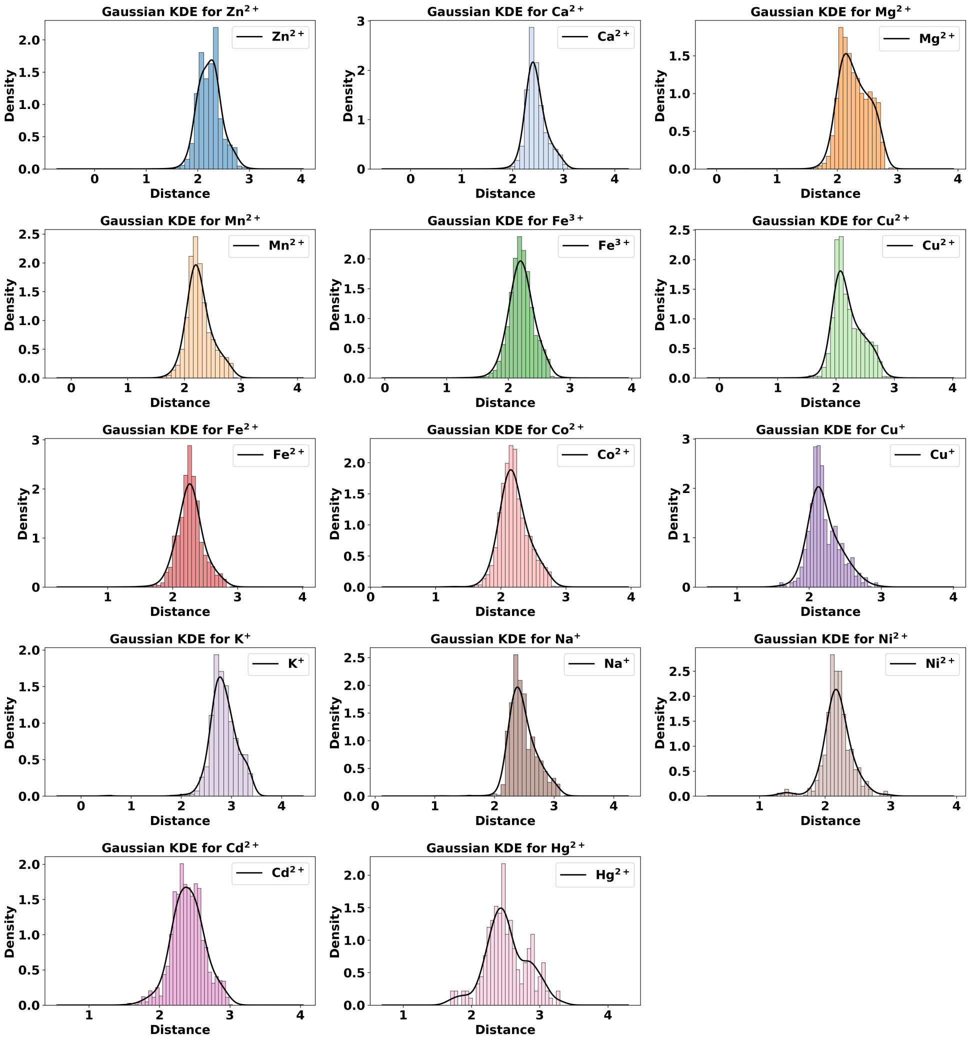


**Fig. S13** The distribution and kernel density estimation (KDE) of distances between binding atoms and metal ions. The distributions exhibit a pronounced clustering tendency across all metal ions.

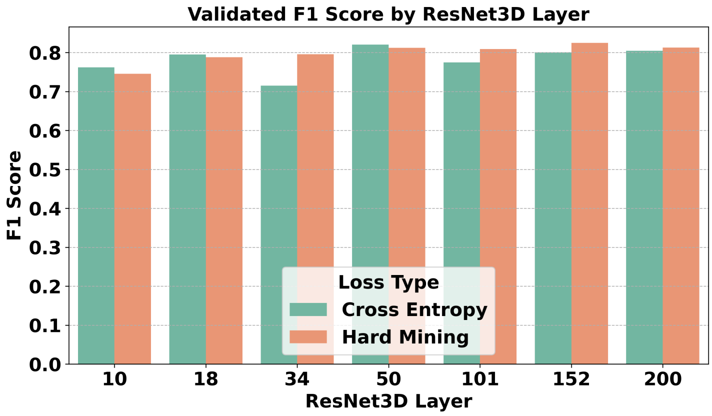


**Fig. S14** Comparison of the cross-entropy loss and the hard mining strategy on the validation set of zinc-binding proteins. The y-axis indicates the validated F1 scores, while the x-axis denotes the number of ResNet3D layers. Green bars represent the validated F1 scores trained with the cross-entropy loss, whereas orange bars depict the scores achieved using the hard mining strategy. The hard mining strategy demonstrates superior and more consistent performance, particularly evident in the two highest bars.

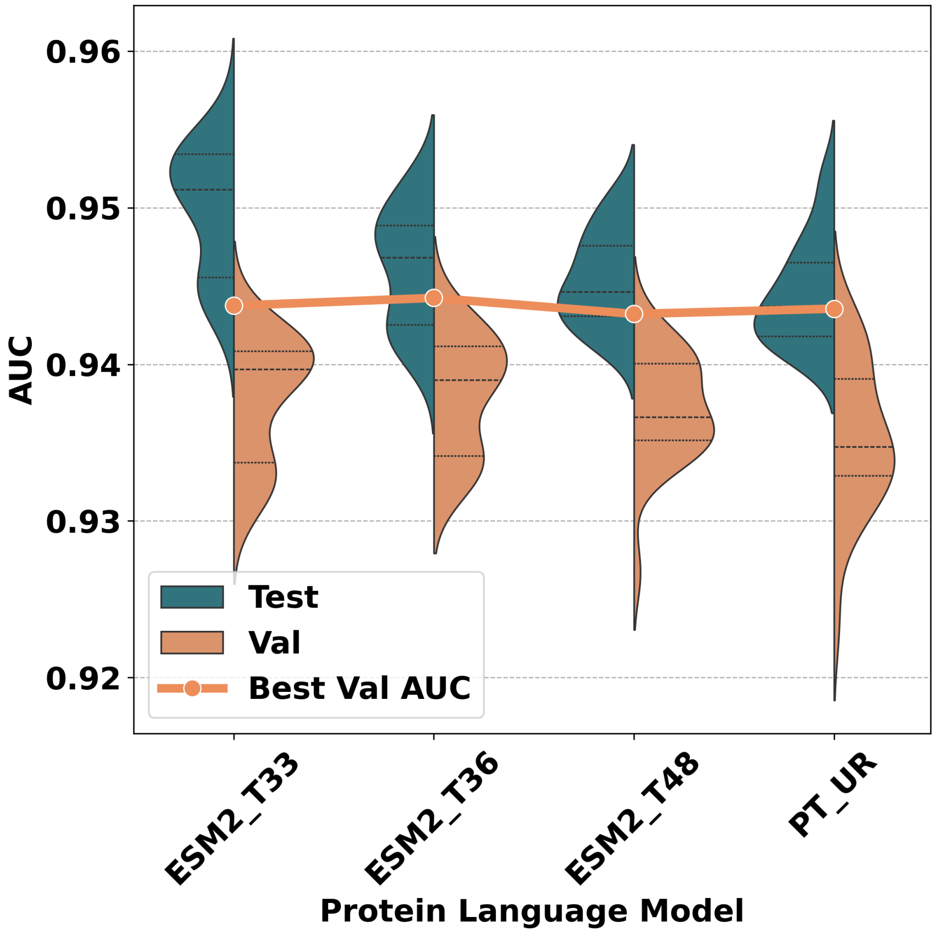


**Fig. S15** Validated and test AUC distributions of PRIME-seq across various protein language models (PLMs). The y-axis represents the AUC scores, while the x-axis denotes the PLMs. Test AUCs (dark green), validated AUCs (orange), and the validated scores of the best models (orange curves) are displayed.

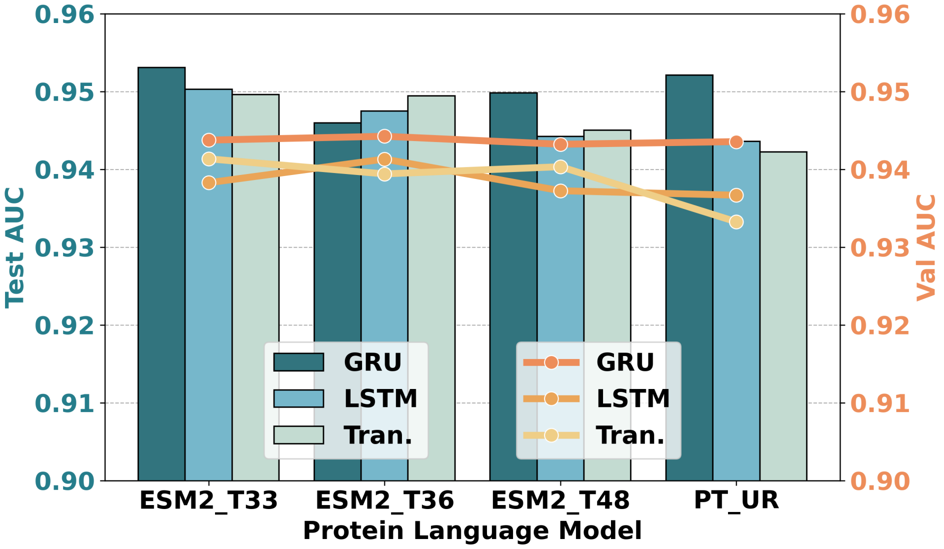


**Fig. S16** Performance of PRIME-seq across various configurations, including interaction layer types and protein language models (PLMs). The left y-axis indicates the test AUC, while the right y-axis denotes the validated AUC. The x-axis categorizes the different PLMs. Interaction layer types are distinguished by color and represented as bars or curves accordingly.

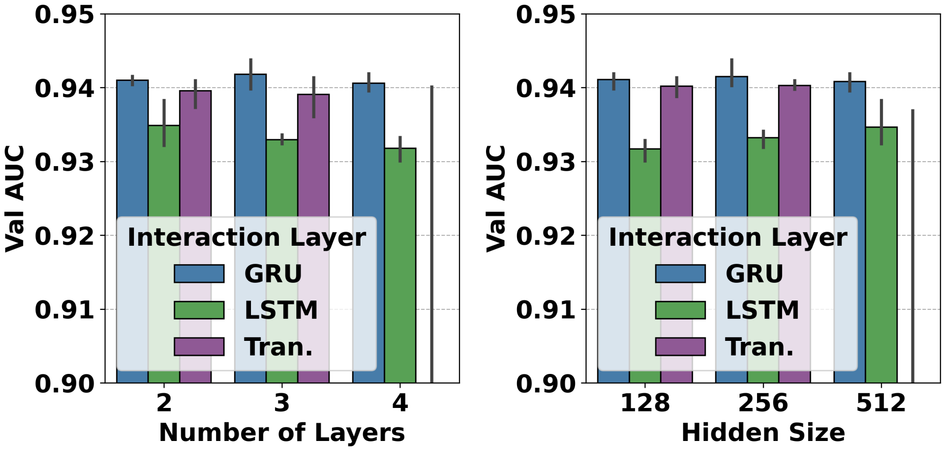


**Fig. S17** Performance of PRIME-seq across various configurations, including interaction layers, hidden sizes, and layer numbers. The y-axis indicates the validated AUC, while the x-axis represents the number of hidden layers (left) and hidden sizes (right). Interaction layer types are distinguished by colors: blue for GRU, green for LSTM, and purple for transformer encoder layers.

### 2 Supplementary tables

**Table S1** Performance of PRIME-seq models on the validation and test sets. This table presents the scores of models computed during the training process, rather than the final evaluation scores, which may exhibit slight variations due to the inherent non-deterministic nature of deep learning models (e.g., dropout, batch normalization, etc.).

| Metal | Val F1 | Test F1 | Val AUC | Test AUC | Val MCC | Test MCC |
|---|---|---|---|---|---|---|
| $Zn^{2+}$ | 0.5291 | 0.7676 | 0.9792 | 0.9904 | 0.5677 | 0.7710 |
| $Ca^{2+}$ | 0.4932 | 0.5494 | 0.9274 | 0.9320 | 0.4916 | 0.5518 |
| $Mg^{2+}$ | 0.2252 | 0.3348 | 0.8822 | 0.9058 | 0.2506 | 0.3692 |
| $Mn^{2+}$ | 0.0506 | 0.0154 | 0.9749 | 0.9543 | 0.1153 | 0.0287 |
| $Fe^{3+}$ | 0.0577 | 0.0390 | 0.9835 | 0.9725 | 0.1051 | 0.0799 |
| $Cu^{2+}$ | 0.2080 | 0.4956 | 0.9379 | 0.9629 | 0.2639 | 0.5280 |
| $Fe^{2+}$ | 0.0000 | 0.0000 | 0.9899 | 0.9954 | 0.0000 | 0.0000 |
| $Co^{2+}$ | 0.0000 | 0.0000 | 0.9383 | 0.9710 | 0.0000 | 0.0000 |
| $Cu^{+}$ | 0.0000 | 0.0000 | 0.9992 | 0.9925 | 0.0000 | 0.0000 |
| $K^{+}$ | 0.0000 | 0.0000 | 0.9469 | 0.9458 | 0.0000 | 0.0000 |
| $Na^{+}$ | 0.4189 | 0.1481 | 0.9538 | 0.7994 | 0.4197 | 0.1497 |
| $Ni^{2+}$ | 0.0000 | 0.0000 | 0.9841 | 0.9931 | 0.0000 | 0.0000 |
| $Cd^{2+}$ | 0.0000 | 0.0000 | 0.9578 | 0.9709 | 0.0000 | 0.0000 |
| $Mn^{3+}$ | 0.7500 | 0.0000 | 1.0000 | 0.9855 | 0.7746 | 0.0000 |
| $Hg^{2+}$ | 0.0000 | 0.0000 | 0.8494 | 0.6479 | 0.0000 | 0.0000 |

**Table S2** Statistical summary of probes for each metal ion following the probe generation process. The table presents the total number of probes (#Probes), the count of positive probes (#Positives), the count of missing probes (#Missing), the number of binding sites (#Sites), the number of proteins (#Proteins), and the mean number of probes per protein (Probes per protein).

| Metal | #Probes ↓ | #Positives ↑ | #Missing ↓ | #Sites | #Proteins | Probes per protein ↓ |
|---|---|---|---|---|---|---|
| $Zn^{2+}$ | 767,872 | 38,961 | 8 | 4,193 | 2,748 | 279 |
| $Ca^{2+}$ | 1,643,317 | 40,601 | 6 | 3,359 | 2,031 | 809 |
| $Mg^{2+}$ | 2,506,839 | 30,873 | 1 | 2,439 | 2,001 | 1,253 |
| $Mn^{2+}$ | 261,795 | 10,036 | 2 | 961 | 670 | 391 |
| $Fe^{3+}$ | 38,719 | 3,970 | 0 | 486 | 368 | 105 |
| $Cu^{2+}$ | 65,770 | 3,114 | 1 | 325 | 225 | 292 |
| $Fe^{2+}$ | 12,612 | 1,521 | 0 | 184 | 156 | 81 |
| $Co^{2+}$ | 142,986 | 3,386 | 1 | 346 | 262 | 546 |
| $Cu^{+}$ | 4,701 | 866 | 0 | 127 | 64 | 73 |
| $K^{+}$ | 85,930 | 1,099 | 0 | 86 | 63 | 1,364 |
| $Na^{+}$ | 80,788 | 1,702 | 0 | 119 | 94 | 859 |
| $Ni^{2+}$ | 8,283 | 619 | 0 | 67 | 55 | 151 |
| $Cd^{2+}$ | 11,728 | 913 | 0 | 109 | 53 | 221 |
| $Hg^{2+}$ | 4,342 | 241 | 0 | 30 | 20 | 217 |

**Table S3** Protein chains containing metal-binding sites omitted by the probee generation algorithm, formatted as "⟨PDB ID⟩⟨chain ID⟩".

| Metal | Missing chains |
|---|---|
| $Zn^{2+}$ | 1r85A, 4tmaE, 5m0tA, 5r4qA, 5zzuB, 6xreA, 7n9zG, 7qg4A |
| $Ca^{2+}$ | 1q8hA, 4ylqL, 7k72A |
| $Mg^{2+}$ | 1vzmA |
| $Mn^{2+}$ | 3rl3A, 6qv4A |
| $Fe^{3+}$ | (None) |
| $Cu^{2+}$ | 6c40B |
| $Fe^{2+}$ | (None) |
| $Co^{2+}$ | 1rxtC |
| $Cu^{+}$ | (None) |
| $K^{+}$ | (None) |
| $Na^{+}$ | (None) |
| $Ni^{2+}$ | (None) |
| $Cd^{2+}$ | (None) |
| $Hg^{2+}$ | (None) |

**Table S4** van der Waals radii of metal ions utilized in the curation of the BioLiP2 datasets.

| Metal | VDW radius |
|---|---|
| C | 1.70 |
| N | 1.55 |
| O | 1.52 |
| S | 1.80 |
| $Zn^{2+}$ | 0.74 |
| $Ca^{2+}$ | 1.00 |
| $Mg^{2+}$ | 0.72 |
| $Mn^{2+}$ | 0.83 |
| $Fe^{3+}$ | 0.645 |
| $Cu^{2+}$ | 0.73 |
| $Fe^{2+}$ | 0.78 |
| $Co^{2+}$ | 0.745 |
| $Na^{+}$ | 1.02 |
| $Cu^{+}$ | 0.77 |
| $K^{+}$ | 1.38 |
| $Ni^{2+}$ | 0.69 |
| $Cd^{2+}$ | 0.95 |
| $Hg^{2+}$ | 1.02 |

[1]The van der Waals radii for the elements in the dataset were derived from the literature [71].

**Table S5** Distribution of metal-binding sites for diverse metal ions across the training, validation, and test datasets.

| Metal | Train | Val | Test |
|---|---|---|---|
| $Zn^{2+}$ | 3,357 | 395 | 441 |
| $Ca^{2+}$ | 2,723 | 272 | 364 |
| $Mg^{2+}$ | 1,991 | 239 | 209 |
| $Mn^{2+}$ | 793 | 90 | 78 |
| $Fe^{3+}$ | 398 | 43 | 45 |
| $Cu^{2+}$ | 274 | 27 | 24 |
| $Fe^{2+}$ | 153 | 15 | 16 |
| $Co^{2+}$ | 277 | 38 | 31 |
| $Cu^{+}$ | 112 | 7 | 8 |
| $K^{+}$ | 68 | 10 | 8 |
| $Na^{+}$ | 93 | 13 | 13 |
| $Ni^{2+}$ | 56 | 6 | 5 |
| $Cd^{2+}$ | 85 | 13 | 11 |
| $Hg^{2+}$ | 24 | 2 | 4 |
| Total | 10,404 | 1,170 | 1,257 |

**Table S6** Number of chains for different metal ions in train/val/test sets.

| Metal | Train | Val | Test |
|---|---|---|---|
| $Zn^{2+}$ | 2,034 | 254 | 255 |
| $Ca^{2+}$ | 1,601 | 200 | 201 |
| $Mg^{2+}$ | 1,736 | 217 | 218 |
| $Mn^{2+}$ | 520 | 65 | 65 |
| $Fe^{3+}$ | 278 | 35 | 35 |
| $Cu^{2+}$ | 156 | 19 | 20 |
| $Fe^{2+}$ | 120 | 15 | 15 |
| $Co^{2+}$ | 216 | 27 | 27 |
| $Cu^{+}$ | 48 | 6 | 7 |
| $K^{+}$ | 50 | 6 | 7 |
| $Na^{+}$ | 74 | 9 | 10 |
| $Ni^{2+}$ | 42 | 5 | 6 |
| $Cd^{2+}$ | 42 | 5 | 6 |
| $Hg^{2+}$ | 16 | 2 | 2 |
| Total | 6,933 | 865 | 874 |

**Table S7** Thresholds for each metal to achieve recall > 0.99 on the validation set. These thresholds are employed in probe detection, wherein a binding residue with a predicted probability exceeding the threshold is identified as a potential binding residue, subsequently verified through structure-based prediction.

| Metal | Threshold for recall > 0.99 |
|---|---|
| $Zn^{2+}$ | 0.00007137 |
| $Ca^{2+}$ | 0.00006270 |
| $Mg^{2+}$ | 0.00004524 |
| $Mn^{2+}$ | 0.00001449 |
| $Fe^{3+}$ | 0.00000678 |
| $Cu^{2+}$ | 0.00005083 |
| $Fe^{2+}$ | 0.00000850 |
| $Co^{2+}$ | 0.00003274 |
| $Na^{+}$ | 0.00001329 |
| $Cu^{+}$ | 0.00001437 |
| $K^{+}$ | 0.00000453 |
| $Ni^{2+}$ | 0.00000292 |
| $Cd^{2+}$ | 0.00002515 |
| $Hg^{2+}$ | 0.00000046 |

**Table S8** Architecture of the ResNet models with varying depths. Conv3d(in_channels, out_channels, kernel_size, stride, padding) represents a 3D convolutional layer, BatchNorm3d(out_channels) denotes a batch normalization layer, LeakyReLU(negative_slope) specifies a leaky ReLU activation function, MaxPool3d(kernel_size, stride, padding) indicates a max pooling layer, and FC(in_features, out_features) refers to a fully connected layer. The notation $\leftarrow n\times$ signifies the number of repetitions of the block.

| Models (#layers) | 10 | 18 | 34 | 50 | 101 | 152 | 200 |
|---|---|---|---|---|---|---|---|
| | Conv3d(4, 64, 7, 2, 3)<br>BatchNorm3d(64)<br>LeakyReLU(0.1)<br>MaxPool3d(3, 2, 1) | $\leftarrow 1\times$ | $\leftarrow 1\times$ | $\leftarrow 1\times$ | $\leftarrow 1\times$ | $\leftarrow 1\times$ | $\leftarrow 1\times$ |
| Layer 1 | Conv3d(64, 64, 3, 1, 1)<br>BatchNorm3d(64)<br>ReLU()<br>Conv3d(64, 64, 3, 1, 1)<br>BatchNorm3d(64) | $\leftarrow 2\times$ | $\leftarrow 3\times$ | $\leftarrow 3\times$ | $\leftarrow 3\times$ | $\leftarrow 3\times$ | $\leftarrow 3\times$ |
| Layer 2 | Conv3d(64, 128, 3, 2, 1)<br>BatchNorm3d(128)<br>ReLU()<br>Conv3d(128, 128, 3, 1, 1)<br>BatchNorm3d(128) | $\leftarrow 2\times$ | $\leftarrow 4\times$ | $\leftarrow 4\times$ | $\leftarrow 4\times$ | $\leftarrow 8\times$ | $\leftarrow 24\times$ |
| Downsample 2 | Conv3d(64, 128, 1, 2, 0)<br>BatchNorm3d(128) | $\leftarrow 1\times$ | $\leftarrow 1\times$ | $\leftarrow 1\times$ | $\leftarrow 1\times$ | $\leftarrow 1\times$ | $\leftarrow 1\times$ |
| Layer 3 | Conv3d(128, 256, 3, 2, 1)<br>BatchNorm3d(256)<br>ReLU()<br>Conv3d(256, 256, 3, 1, 1)<br>BatchNorm3d(256) | $\leftarrow 2\times$ | $\leftarrow 6\times$ | $\leftarrow 6\times$ | $\leftarrow 23\times$ | $\leftarrow 36\times$ | $\leftarrow 36\times$ |
| Downsample 3 | Conv3d(128, 256, 1, 2, 0)<br>BatchNorm3d(256) | $\leftarrow 1\times$ | $\leftarrow 1\times$ | $\leftarrow 1\times$ | $\leftarrow 1\times$ | $\leftarrow 1\times$ | $\leftarrow 1\times$ |
| Layer 4 | Conv3d(256, 512, 3, 2, 1)<br>BatchNorm3d(512)<br>ReLU()<br>Conv3d(512, 512, 3, 1, 1)<br>BatchNorm3d(512) | $\leftarrow 2\times$ | $\leftarrow 3\times$ | $\leftarrow 3\times$ | $\leftarrow 3\times$ | $\leftarrow 3\times$ | $\leftarrow 3\times$ |
| Downsample 4 | Conv3d(256, 512, 1, 2, 0)<br>BatchNorm3d(512) | $\leftarrow 1\times$ | $\leftarrow 1\times$ | $\leftarrow 1\times$ | $\leftarrow 1\times$ | $\leftarrow 1\times$ | $\leftarrow 1\times$ |
| | | | AdaptiveAvgPool3d(1)<br>FC(512,out_features) | | | | |

**Table S9** Evaluation of various ResNet3D architectures, pretrained configurations, and loss functions on the zinc-binding validation dataset, presented in terms of true positives (TP), false negatives (FN), false positives (FP), F1 score, and root mean square error (RMSE).

| ResNet3D Layer | Pretrained | Loss | TP | FN | FP | F1 | RMSE |
|---|---|---|---|---|---|---|---|
| 152 | True | hard_mining | 316 | 46 | 88 | 0.8251 | 1.3053 |
| 50 | True | ce | 316 | 54 | 84 | 0.8208 | 1.3424 |
| 200 | True | hard_mining | 309 | 55 | 87 | 0.8132 | 1.2245 |
| 50 | True | hard_mining | 316 | 47 | 99 | 0.8123 | 1.3585 |
| 101 | True | hard_mining | 320 | 45 | 106 | 0.8091 | 1.4317 |
| 200 | True | ce | 309 | 56 | 94 | 0.8047 | 1.2936 |
| 152 | True | ce | 294 | 72 | 74 | 0.8011 | 1.4070 |
| 34 | True | hard_mining | 310 | 52 | 107 | 0.7959 | 1.3084 |
| 18 | True | ce | 293 | 70 | 81 | 0.7951 | 1.2918 |
| 18 | True | hard_mining | 302 | 60 | 102 | 0.7885 | 1.3399 |
| 50 | False | hard_mining | 293 | 71 | 87 | 0.7876 | 1.2951 |
| 34 | False | hard_mining | 286 | 85 | 74 | 0.7825 | 1.3682 |
| 18 | False | ce | 289 | 75 | 91 | 0.7769 | 1.3310 |
| 101 | False | ce | 277 | 91 | 70 | 0.7748 | 1.2716 |
| 50 | False | ce | 299 | 65 | 119 | 0.7647 | 1.3273 |
| 200 | False | hard_mining | 274 | 97 | 74 | 0.7622 | 1.3232 |
| 10 | True | ce | 298 | 72 | 114 | 0.7621 | 1.4757 |
| 18 | False | hard_mining | 280 | 89 | 92 | 0.7557 | 1.2709 |
| 101 | True | ce | 321 | 44 | 172 | 0.7483 | 1.2775 |
| 10 | False | hard_mining | 277 | 90 | 99 | 0.7456 | 1.3747 |
| 10 | False | ce | 296 | 72 | 144 | 0.7327 | 1.4878 |
| 34 | False | ce | 255 | 117 | 86 | 0.7153 | 1.2810 |
| 10 | True | hard_mining | 320 | 45 | 266 | 0.6730 | 1.3760 |
| 34 | True | ce | 321 | 46 | 392 | 0.5944 | 1.3535 |
| 152 | False | hard_mining | 281 | 84 | 381 | 0.5472 | 1.7328 |
| 200 | False | ce | 190 | 196 | 175 | 0.5060 | 1.7025 |
| 101 | False | hard_mining | 316 | 51 | 644 | 0.4763 | 1.5489 |
| 152 | False | ce | 299 | 74 | 1,074 | 0.3425 | 2.0319 |

**Table S10** Optimal validation AUCs achieved by each protein language model (PLM) alongside their respective hyperparameters.

| PLM | AUC | Num Layers | Hidden Size | Interaction Layer |
|---|---|---|---|---|
| ESM2_T33 | 0.943743 | 3 | 256 | GRU |
| ESM2_T36 | 0.944240 | 2 | 256 | GRU |
| ESM2_T48 | 0.943214 | 2 | 512 | GRU |
| PT_UR | 0.943551 | 2 | 512 | GRU |

#### 3 Supplementary Algorithms

To construct a non-redundant dataset for each metal ion, sequences are clustered using `MMseqs2 easy-cluster`. The process begins with the least abundant metal ion, $Hg^{2+}$, and proceeds sequentially to the most abundant, $Zn^{2+}$.

---

**Algorithm S1** Gradual Construction of Non-Redundant Datasets for Metal Ions

---

**Require:** Sequence list $S$ and metal ion list $M$
**Ensure:** Non-redundant sequences $C = \{C_m\}_{m \in M}$

**function** CONSTRUCT_NONREDUNDANT_DATASET$(S, M)$
    Sort $S$ by ascending metal counts
    $C \leftarrow \{\emptyset\}$
    **for** each metal $m$ in $M$ **do**
        $M' \leftarrow \{s \in S | s \text{ contains metal } m\}$
        $M' \leftarrow M' \cup$ `flatten`$(C)$    ▷ *Flatten the list of list*
        Cluster $M'$ into $M''$ with `MMseqs2 easy-cluster`
        $C_m \leftarrow \emptyset$
        **for** each cluster $c$ in $M''$ **do**
            **if** $c \cap$ `flatten`$(C) = \emptyset$ **then**    ▷ *Check redundancy*
                $C_m \leftarrow C_m \cup$ `representative`$(c)$
            **end if**
        **end for**
        $C \leftarrow C \cup \{C_m\}$
    **end for**
    **return** $C$
**end function**

---

**Algorithm S2** Generate Probes for Metal-Binding Sites from Protein Sequences

**Require:** Metal ion type $metal$, all residues $R$, predicted probabilities $Pr$, $\delta \leftarrow$ a globally stored mapping from (residue, atom, metal) to distance thresholds, $t$ the threshold for selected binding residues as listed in Supplementary Table S7

**Ensure:** List of probe positions

**function** GET_PROBE_POSITIONS($metal$, R, $Pr$)
    $P \leftarrow$ coordinates of all N, O, and S atoms in $R$
    $L \leftarrow$ names of all N, O, and S atoms in $R$
    $P' \leftarrow \{d(r) \mid r \in R\}$ ▷ *$d(r)$: grid of N/O/S atoms in residue $r$, with a 5 Å margin and 0.5 Å spacing*
    $D \leftarrow$ Euclidean distance matrix between all points in $P'$ and $P$
    $I \leftarrow$ COLLECT_BINDING_PROBES$(D, L, metal, \delta)$
    Sort $I$ in descending order of $\max Pr[i]$ for each cluster $i$ in $I$
    $Probes \leftarrow \emptyset$
    **for** each cluster $c$ in $I$ **do**
        $pos \leftarrow \{p_i \mid i \in c\}$ ▷ *$p_i$: position of atom $i$ in cluster $c$*
        $p_1 \leftarrow \max_{i \in c} Pr[\text{res}(i)]$ ▷ *res$(i)$: residue index for atom $i$*
        $p_2 \leftarrow \max_{i \in c} Pr[i]$ ▷ *Maximum predicted probabilities*
        **if** $p_1 < 0.001$ or $p_2 < 0.001$ **then**
            **continue** ▷ *Skip if low appearance in dataset*
        **end if**
        $p^* \leftarrow$ center of $pos$ ▷ *Representative probe*
        **if** $p^*$ is far from any probe in $Probes$ **then** ▷ *Check overlapping*
            $Probes \leftarrow Probes \cup \{p^*\}$
        **end if**
    **end for**
    **return** $Probes$
**end function**

**Algorithm S3** Collect Binding Probe Indices Based on Distance Matrix and Atom Names

**Require:** Distance matrix $D \in \mathbb{R}^{N \times M}$, atom names $L \in \mathbb{R}^{M}$, metal ion type $m$, distance thresholds $\delta$

**Ensure:** List of filtered probes

**function** COLLECT_BINDING_PROBES$(D, L, m, \delta)$
  $N, M \leftarrow$ shape of $D$ ▷*Number of probes and atoms*
  Initialize $\Delta$ as zero Boolean tensor of shape $(M, 2)$
  **for** each $i$ in 1 to $M$ **do**
    $l, u \leftarrow \delta[L[i], m]$ ▷*Lower and upper bounds of distance between atom name $L[i]$ and metal $m$*
    $\Delta[L[i], 0] \leftarrow l$
    $\Delta[L[i], 1] \leftarrow u$
  **end for**
  $\delta_{\min} \leftarrow \min_x \Delta[x, 0]$
  $D' \leftarrow (D > \delta_{\min})\ \&\ (D > \Delta[:, 0])\ \&\ (D < \Delta[:, 1])$ ▷*Bitwise operations*
  $D' \leftarrow$ PACK_BOOLEAN_TENSOR$(D')$ ▷*For efficiency. $D' \in \mathbb{N}^{N \times \lceil M/8 \rceil}$*
  Cluster $D'$ into $C$ accroding to its rows ▷*For $i, j \in \{0, 1, \dots, N-1\}$, $D'[i] = D'[j]$ means $i$ and $j$ are in the same cluster.*
  **return** $C$
**end function**

**Algorithm S4** Efficient Packing of Boolean Tensor into Byte Tensor

**Require:** $x \in \{0, 1\}^{B \times N}$ ▷*Boolean tensor*

**Ensure:** $output \in \mathbb{N}^{B \times \lceil N/8 \rceil}$ ▷*Packed byte tensor*

**function** PACK_BOOLEAN_TENSOR$(x)$
  $(B, N) \leftarrow$ shape of $x$
  $M \leftarrow \lceil N/8 \rceil$
  Initialize *output* as zero tensor of shape $(B, M)$ ▷`dtype=torch.uint8`
  **for** $i = 0$ to $N - 1$ **do**
    $j \leftarrow i/8$
    $k \leftarrow i \bmod 8$
    $output[:, j] \leftarrow output[:, j] \mid (x[:, i] \ll k)$ ▷*Vectorized operations*
  **end for**
  **return** *output*
**end function**

**Algorithm S5** Efficient Unpacking of Boolean Tensor from Byte Tensor

**Require:** $x \in \mathbb{N}^{B \times M}$, $N \in \mathbb{N}$ ▷ *Byte tensor and number of bits*
**Ensure:** $output \in \{0, 1\}^{B \times N}$ ▷ *Unpacked boolean tensor*

**function** UNPACK_BOOLEAN_TENSOR$(x, N)$
  assert $N \leq 8 \times M$ ▷ *Ensure N is within bounds*
  $(B, M) \leftarrow$ shape of $x$
  Initialize *output* as zero tensor of shape $(B, N)$ ▷ `dtype=torch.bool`
  **for** $i = 0$ to $N - 1$ **do**
    $j \leftarrow i/8$
    $k \leftarrow i \bmod 8$
    $output[:, i] \leftarrow (x[:, j] \gg k)\ \&\ 1$ ▷ *Vectorized operations*
  **end for**
  **return** *output*
**end function**

**Algorithm S6** Postprocessing of Probes for Metal-Binding Site Prediction

**Require:** $P \subset \mathbb{R}^3$ ▷ *List of probe positions predicted as binding sites*
**Ensure:** $P^* \subset \mathbb{R}^3$ ▷ *Output probe positions*

**function** POSTPROCESS_PROBES$(P)$
  Cluster $P$ into $C$ with `scipy.cluster.hierarchy.fclusterdata` ▷ *Cluster probes within 3.0 Å or customized distance*
  $H \leftarrow \emptyset$
  **for** each cluster $c$ in $C$ **do**
    Choose $p^* \in c$ with the highest probe probability
    $H \leftarrow H \cup \{(p^*, prob, |c|)\}$ ▷ *Store probe position, probability, and cluster size*
  **end for**
  $P^* \leftarrow \emptyset$
  **for** each $p*, prob, |c|$ in $H$ **do**
    **if** $prob \geq 0.5$ or $|c| \geq 2$ **then** ▷ *Default 0.5 threshold, can be customized*
      $P^* \leftarrow P^* \cup \{p^*\}$
    **end if**
  **end for**
  **return** $P^*$
**end function**
